# Spatial multi-omics resolve epithelium-fibroblast gradients and highlight NESTIN-NOTCH1-expressing subepithelial fibroblasts during human pancreatic tumorigenesis

**DOI:** 10.64898/2026.09.04.749182

**Authors:** Maëlle Batardière, Myriam Iliana Ibanez-Rios, Abdelhakim Khellaf, Elham Dianati, Sarah-Slim Diwan, Maxence Pelloux, Alexandre Archambault-Marsan, Camille Beaussier, Jade Diwan, Sasha Sapon-Cousineau, Jumanah Baig, Ayman Shoukari, Zean Ghanmeh, Zhiyuan Yang, Melisa Farias Gonzalez, Jia-Lin Li, Ali Kassab, Leonardo Lando, Philippe Lefrancois, Simon Turcotte, Youngmin A Lee, Simon F Roy, Mahdi S Hosseini, Basile Tessier-Cloutier, Marcus CB Tan, Kathleen E DelGiorno, David JHF Knapp, Vincent Q Trinh

**Author notes:** **Corresponding author:** Quoc-Huy Vincent Trinh, MD MSc FRCPC, Institute for Research in Immunology and Cancer, 2950 chemin de la Polytechnique, Room 3440, Montréal, QC, H3T 1J4, Canada.

## Abstract

Intraductal papillary mucinous neoplasms (IPMNs) are cystic precursors of pancreatic ductal adenocarcinoma undergoing dynamic epithelial and stromal remodeling during progression. In this study, we integrated computational pathology, cyclic immunofluorescence, and Xenium spatial transcriptomics to define the spatial organization of the IPMN microenvironment. We queried cell-based histopathological features and identified stromal patterns strongly associated with epithelium proximity. Cyclic immunofluorescence and Xenium revealed a subepithelial gradient extending from myofibroblasts to inflammatory fibroblasts. Xenium further identified a distinct subepithelial *NOTCH1*-*NESTIN*-expressing fibroblast subset in myofibroblasts that expands with tumor progression. By recapitulating the spatial remodeling event using human pancreatic tumor cells and primary myofibroblast co-culture models, we demonstrated the capacity of pancreatic tumor cells to induce a NESTIN-expressing state in neighboring myofibroblasts. These findings reveal new dynamic epithelial-stromal interactions through subepithelial NESTIN+ fibroblasts associated with pancreatic tumorigenesis and immediate microenvironment remodeling by active tumor cells, highlighting the power of computational pathology integrated with spatial transcriptomics.

**STATEMENT OF SIGNIFICANCE:** By integrating computational pathology, cyclic immunofluorescence, and Xenium spatial transcriptomics, we show that epithelium proximity drives fibroblast identity in IPMNs. We reveal the existence of subepithelial NESTIN-expressing fibroblasts associated with disease progression, and the capacity of human pancreatic tumor cells to remodel myofibroblasts in close contact into NESTIN-expressing fibroblasts.

## INTRODUCTION

Pancreatic ductal adenocarcinoma (PDAC) is one of the deadliest solid malignancies, with a 5-year survival below 13% (1) and a global mortality burden that ranks it among the leading causes of cancer-related death worldwide (2). Fewer than 20% are eligible for resection because most patients are diagnosed at an advanced stage. The greatest opportunity to alter the course of PDAC lies upstream of clinical presentation. PDAC arises predominantly from two precursor lesions: microscopic pancreatic intraepithelial neoplasia (PanIN) and macroscopic, cystic intraductal papillary mucinous neoplasms (IPMNs) (3). IPMNs are readily detected on routine imaging and, in principle, are resectable before invasive transformation (4). Both precursors undergo a transition from low-grade (LG) to high-grade (HG) to invasive cancer (INV). Yet this window is poorly exploited: management remains binary, alternating between surveillance and surgery with no non-invasive means to halt progression (5). A more granular understanding of the events governing the transition from low-grade precursor to invasive carcinoma is needed to identify new therapeutic opportunities.

Most studies on the transition from pancreatic precursors to invasive PDAC focus on genomic and epigenetic alterations in the epithelial component (6). Advances in single-cell transcriptomics and spatial technologies have enabled a deeper understanding of IPMN molecular pathogenesis. Precursor is accompanied by pronounced heterogeneity within the neoplastic epithelium, and specific gene expression profiles were associated with histological progression, providing a basis for defining high-risk populations as indicators for early intervention (7–9). Single-cell and lineage analyses have resolved a spectrum of epithelial states, including stem-like or progenitor-like populations implicated in seeding and sustaining progression (3, 10, 11). We have recently published data showing the presence of pyloric metaplasia features with CD44v9 and AQP5 expression increasing with IPMN progression in human samples (12).

The tumor microenvironment (TME) is a recognized driver of progression in multiple cancers (13–16). It is believed to be particularly consequential in pancreatic disease due to its characteristically dense stroma (17–19). In PDAC, the TME is complex and dominated by cancer-associated fibroblasts (CAFs), which have been broadly resolved into recurrent subtypes, including myofibroblastic (myCAF), inflammatory (iCAF), and antigen-presenting (apCAF) populations (20). Specifically, myCAFs are TGFβ-driven, αSMA^HIGH^, and responsible for ECM deposition. iCAFs are αSMA^LOW^, driven by IL-1/JAK-STAT signaling, and immunosuppressive (20, 21). A growing body of work reframes these categories as interconvertible states along a continuum rather than fixed lineages, shaped by plasticity and local context (22). CAFs were first described as solely tumor-promoting, but are in reality functionally heterogeneous and can harbor tumor-restraining phenotypes (23). Depletion of αSMA+ CAFs increases tumor aggressiveness (24), while IL1/JAK signaling inhibition, increasing myCAF to iCAF ratio, resulted in better prognosis (23). Critically, at the invasive stage, fibroblast organization is spatially structured rather than random, with current dogma describing myCAFs as juxtatumoral, while iCAFs reside more distally, therefore presenting identities driven by tumor distance (25).

As with PDAC, IPMN lesions are surrounded by a dense and fibrotic stroma, but less is known about CAF heterogeneity and function during the pre-invasive stage. Single-cell RNAseq studies have reported the presence of myCAF, iCAF, and apCAF in HG-IPMN, while only myCAF were rarely detected in LG-IPMN (26, 27). Spatially resolved studies of IPMN have largely focused on the epithelial compartment and the immune infiltrate (7, 8, 28, 29), but one study briefly highlights enrichment of myCAF concomitant with HG-IPMN to PDAC progression (8). The spatial behavior of fibroblasts relative to distance from the evolving epithelium remains uninvestigated.

Our study set out to define how the stromal microenvironment is spatially organized relative to the neoplastic epithelium across the stages of pancreatic tumorigenesis. We leveraged IPMN heterogeneity to reconstruct the LG, HG, and INV sequence and focused on stromal visual cues using computational pathology methods combined with spatial statistics. Subepithelial stromal variations defined disease progression. We queried these patterns with cyclic immunofluorescence and homed in on potential signaling pathways with spatial transcriptomics combined with novel distance-based methods. We identified the NOTCH1-NESTIN pathway as a prevalent tumor-induced subepithelial component in high-grade precursors. We validated the capacity of tumor cells to remodel their immediate stromal microenvironment *in vitro*, with mechanistic validation in a co-submission (Ibanez-Rios *et al*.). As we show that tumor cells readily modulate their immediate stromal microenvironment, our results nuance concepts of defined fibroblast functional classes and layers of specific fibroblast phenotypes.

## RESULTS

### Computational pathology analysis of IPMN stroma highlights fibroblast-rich changes in proximity to epithelial cells during disease progression

To investigate the evolution of histomorphological architecture during pancreatic tumorigenesis, we built a computational pathology pipeline for cell-centered feature extraction using UNI2-h and unsupervised clustering of features to reveal main cellular patterns across the disease landscape (**Fig. 1A**). This method uses 20X magnification whole-slide images from hematoxylin and eosin stains. This approach queried regions of interest from 142 normal ducts, 146 LG, 84 HG, and 28 INV IPMN from 83 patients (52.4% female) targeted by two board-certified pathologists. We extracted 777,710 cell-centered patches: 51,985 from normal duct areas, 358,232 LG, 269,121 HG, and 98,372 INV. Feature extraction with UNI2-h followed by K-means (k=35) clustering of the embeddings was performed (**Fig. 1B**). Three board-certified pathologists reviewed representative cell-centered tiles, confirmed consistent grouping of cellular histopathological patterns, and provided semantic characterization of each cluster, including whether it contained epithelial or stromal components (**Supplementary Fig. S1A**). Analysis of cluster proportions by grade confirmed a shift in stromal histomorphological architecture with disease progression (**Fig. 1C**, **Supplementary Fig. S1B-D**).

**Figure 1.**
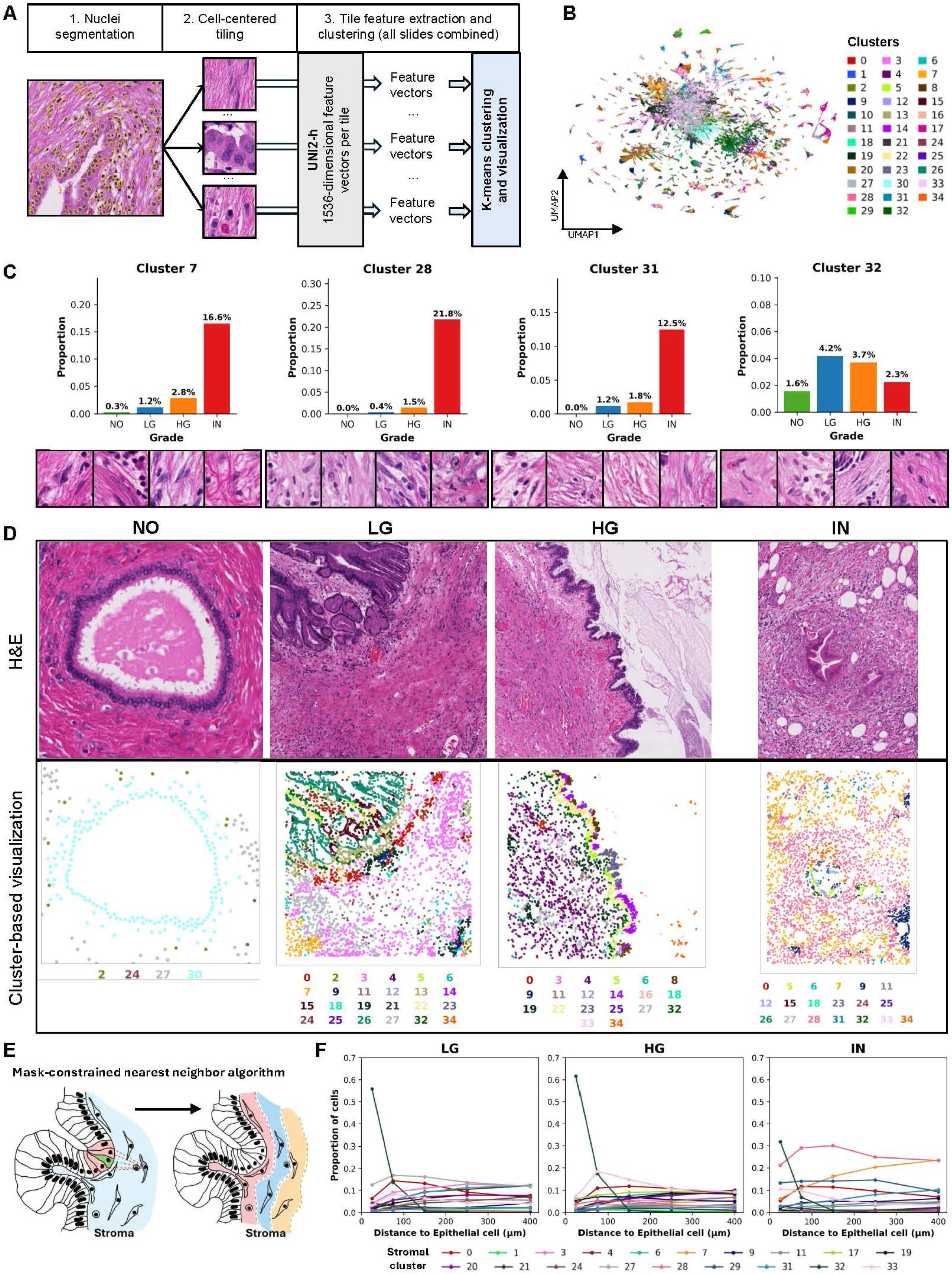
Computational pathology analysis of IPMN stroma highlights fibroblast-rich changes in proximity to epithelial cells during disease progression. **A**. Schematic overview of the cell-centered clustering computational pathology processing pipeline **B.** UMAP projection of cell-centered patches colored per K-mean clustering results run on UNI2 extracted features. **C.** Bar plot of proportion of patch cluster per grade, and representative patch cluster content (7, 28, 31, and 32) **D.** H&E images of N, LG, HG and INV regions with associated representative projection of cell-centered patches centroids colored per cluster. **E.** Schematic overview of the stroma focused nearest neighbour algorithm. **F**. Plot of proportions of stromal patches clusters as a function of distance to nearest epithelial patch.

More specifically, we observed gradual, grade-dependent changes in epithelial cluster proportions, with semantic annotation of grade-specific atypia or benign cluster state matching what clustering alone reveals (**Supplementary Fig. S1A-B**). Alongside epithelial evolution, our data showed concomitant stromal remodeling with disease progression. Notable stromal clusters include desmoplastic stroma with occasional plasmocytic infiltration (cluster 7), desmoplastic stroma with mixed inflammatory cells (cluster 28), desmoplastic stroma with occasional myofibroblastic features (cluster 31), and subepithelial stroma with fibroblasts, collagen, capillaries, and occasional immune cells (cluster 32) (**Fig. 1C**). These clusters were nearly non-existent in normal tissue, gradually increased in LG and HG, and became the most prominent fraction of invasive stroma (**Fig. 1C** and **Supplementary Fig. S1B-D**).

Spatial projection of cluster-assigned patches revealed gradients deriving from the epithelial clusters (**Fig. 1D**). Subepithelial areas (100 µm) showed the most significant shifts with disease progression (**Fig. 1D**). To statistically validate these findings, we developed a mask-constrained nearest neighbor algorithm to identify the closest epithelial component and its distance for each stromal patch (**Fig. 1E**). Notably, cluster 32 was largely prominent in the subepithelial stroma of all LG, HG, and INV grades, within approximately 50 µm. In INV, cluster 28 was observed in both proximal and distant stroma, while clusters 7 and 31 were enriched distally (**Fig. 1F**). Overall, we reveal heterogeneous stromal remodeling during pancreatic tumorigenesis, with fibroblast-rich stromal patterns shifting relative to tumor proximity, particularly in the subepithelial region.

### CyCIF of IPMN samples highlight epithelial stem cell signatures and fibroblast lineage alterations during tumor progression

To quantify epithelial and stromal cellular heterogeneity at single-cell resolution during pancreatic tumorigenesis, we deployed cyclic immunofluorescence (CyCIF) on 16 whole-slide tissue sections from FFPE human samples (**Fig**. **2A**, **Supplementary Table 1**). We designed a panel of 19 markers targeting the diversity of known epithelial (CD44v9, CD166, CD133, S100P, SPP1), stromal (CXCL12, COL1A1, CD74, CD105, αSMA, FAP), immune (CD45, CD4, CD8, CD11b, CD3, CD163, CD20), and endothelial (CD31) markers (**Fig. 2B** and **Supplementary Table 2**). V.Q.T. performed grade-specific region of interest (ROI) tissue annotations, generating 20 regions of interest of normal ducts, 9 of acinar-to-ductal metaplasia/chronic pancreatitis (ADM), 73 of low-grade IPMN, 31 of high-grade IPMN, and 23 of PDAC, each centered on the epithelial-to-stromal interface (**Supplementary Table 2**). Stromal and epithelial cells are the most prevalent populations within the queried areas, with no strong trends in cell composition across disease progression (**Fig. 2C-E**). To decode epithelial phenotypic heterogeneity, we clustered epithelial cells based on the expression of epithelial-specific markers (CD44v9, CD166, CD133, S100P, SPP1). We identified 7 subpopulations, annotated by marker expression profiles, that varied with disease progression (**Fig. 2F-I**). Similarly, we clustered stromal subpopulations based on stromal-specific markers (CXCL12, COL1A1, CD74, CD105, αSMA, FAP) and highlighted 10 subpopulations that also varied with disease progression (**Fig. 2J-M**). Overall, we observed variations within both the epithelial and stromal compartments with IPMN progression.

**Figure 2.**
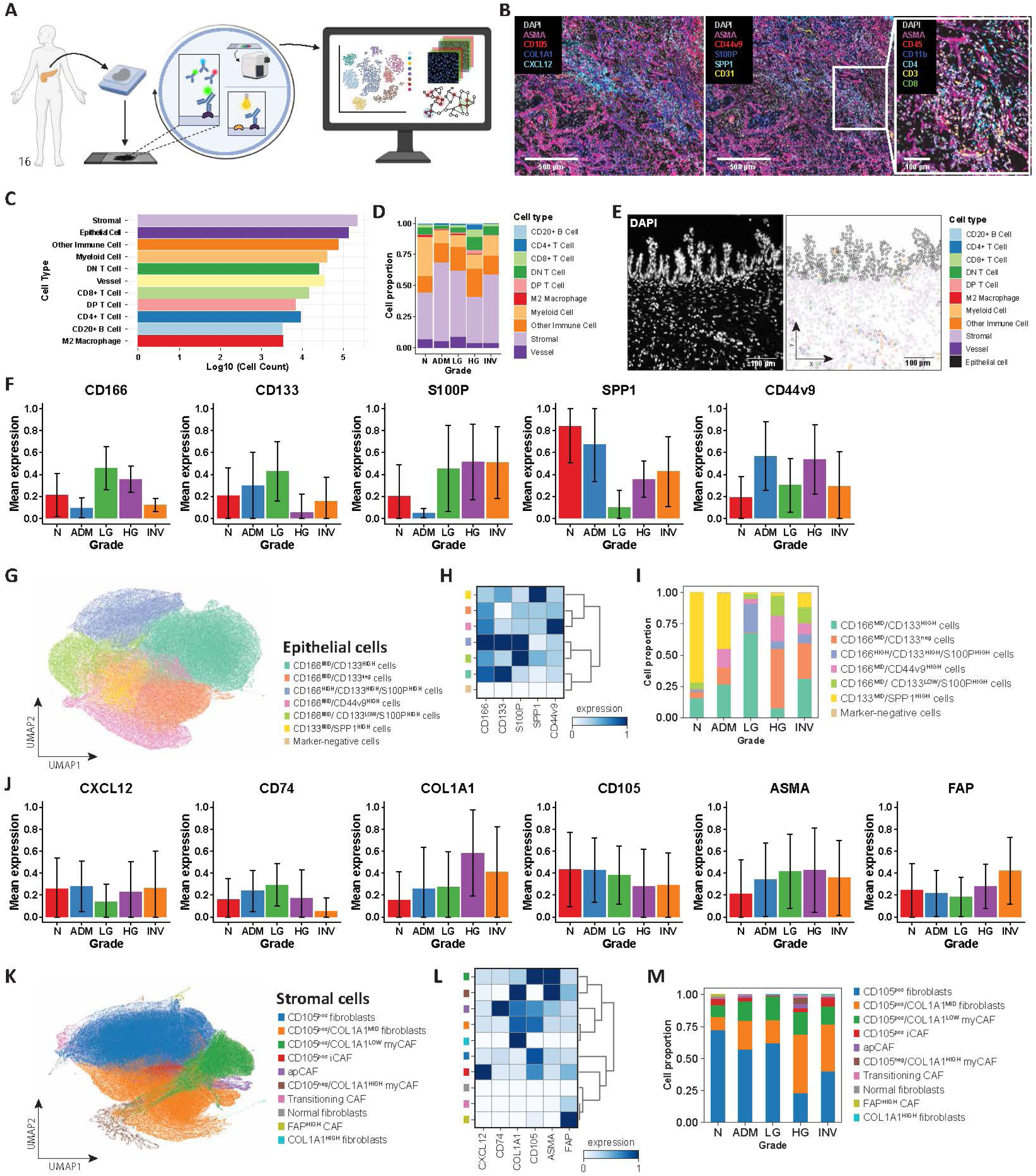
CyCIF of IPMN samples highlight epithelial stem cell signatures and fibroblast lineage alterations during tumor progression. **A.** Schematic overview of the experimental design for multiplexed cyclic immunofluorescence. Created in Biorender. **B.** Representative image of invasive PDAC showing DAPI, ASMA, CD105, COL1A1, CXCL12; DAPI, ASMA, CD44v9, S100P, SPP1, CD31, and DAPI, ASMA, CD45, CD11b, CD4, CD3, CD8 signals respectively. **C**. Bar plot of cell counts per cell type (Log 10). **D**. Stacked bar graph of cell type proportions per grade, excluding epithelial cells. **E**. Representative high-grade dysplasia tissue sections showing DAPI signal (left) and localization of cells colored by cell type (right). **F**. Histogram of the average expression of the markers CD166, CD133, S100P, SPP1, and CD44v9 by each epithelial cell according to grade. **G**. UMAP projection of single epithelial cells, from n=16 samples, with color-coded Leiden clusters annotated according to expression profiles. **H**. Matrix plot of expression levels of CD166, CD133, S100P, SPP1, and CD44v9 per Leiden cluster color-coded as in (**G**). **I**. Stacked bar graph of cell proportion of each epithelial cell cluster per grade. **J**. Histogram of the average expression of the markers CXCL12, CD74, COL1A1, CD105, αSMA, and FAP by each stromal cell according to grade. **K**. UMAP projection of single stromal cells, from n=16 samples, with color-coded Leiden clusters annotated according to expression profiles. **L**. Matrix plot of expression levels of CXCL12, CD74, COL1A1, CD105, αSMA, and FAP per Leiden cluster color-coded as in (**K**). **M**. Stacked bar graph of cell proportion of each stromal cell cluster per grade. N - Normal, ADM - acinar-to-ductal metaplasia, LG - low-grade dysplasia, HG - High-grade dysplasia, INV - invasive cancer.

### Epithelial-stromal distance shapes fibroblast identity

The observed variations do not account for spatial distributions, notably in the precursor stage, where epithelial cells form a lining against the stroma. As the previous computational pathology analysis suggested gradients emerging from the epithelial component, we performed the same mask-constrained epithelial-stromal analysis with our imaging data. Several stromal markers showed spatial biases, including αSMA enriched in epithelial-adjacent stroma and COL1A1 in distal stroma (**Fig. 3A**). To quantify this bias, we applied our mask-constrained nearest-neighbor algorithm, assigning each stromal cell its distance to the nearest epithelial cell. Grade-specific UMAP embeddings of single stromal cells, colored by nearest distance to epithelial cells, revealed a spatial organization that evolved with grade (**Fig. 3B**). At the LG stage, cells formed a fragmented embedding, with epithelial-proximal cells occupying separate clusters from epithelial-distal cells. At the HG stage, the embedding was continuous and cohesive, with epithelial-proximal cells concentrated superiorly and a smooth gradient toward epithelial-distal cells in the inferior region of the projection. This pattern was less observed in INV disease.

**Figure 3.**
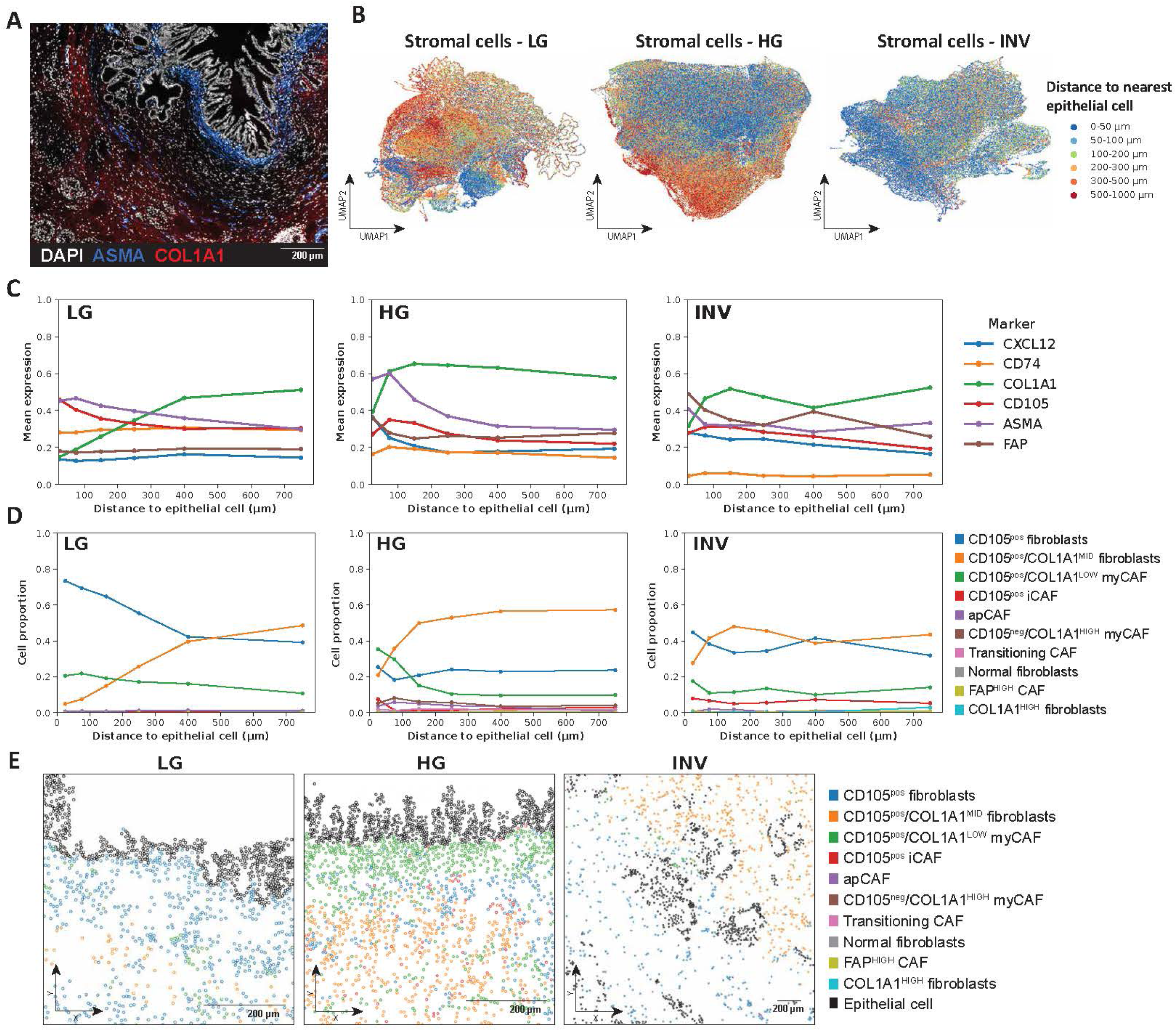
Epithelial-stromal distance shapes fibroblast identity. **A**. Representative image of high-grade dysplasia showing signal for DAPI (in white), ASMA (in blue), and COL1A1 (in red). **B**. UMAP projection of single stromal cells, from n=16 samples, color-coded according to their distance to the nearest epithelial cell, for respectively LG, HG, and INV cells only. **C**. Connected dot plot of expression levels of markers CXCL12, CD74, COL1A1, CD105, αSMA, and FAP as a function of stromal cell distance to nearest epithelial cell. **D**. Connected dot plot of proportions of stromal sub-populations as a function of stromal cell distance to nearest epithelial cell. **E**. Representative tissue sections showing the localization of epithelial cells (in black) and stromal cell populations of LG, HG, and INV regions. Stromal cells are colored according to Leiden clustering results. LG - low-grade dysplasia, HG - High-grade dysplasia, INV - invasive cancer. Distance ranges for connected line plots: 0-50 μm, 50-100 μm, 100-200 μm, 200-300 μm, 300-500 μm, 500-1000 μm.

To identify which markers were associated with spatial distribution relative to the epithelium, we examined marker expression levels across increasing distance from epithelial cells (**Fig. 3C**). At low-grade, ASMA and CD105 levels decreased slightly with increasing distance to the epithelium, whereas COL1A1 levels increased with distance, and CXCL12, CD74, and FAP remained stable. The transition to high-grade revealed a steeper decrease in ASMA outside the immediate peri-epithelial niche, and at the invasive stage, the decrease with distance was smaller. COL1A1 levels remained higher in the distal stroma, at both high-grade and invasive stages, whereas FAP was elevated near PDAC tumor cells. Analysis of stromal subpopulation proportions relative to epithelium distance in low-grade showed that the peri-epithelial region was dominated by CD105^pos^ fibroblasts, while CD105^pos^/COL1A1^MID^ fibroblasts predominated in regions further away from the epithelium (**Fig. 3D-E**). We observed a shift at high-grade, with enrichment of CD105^pos^/COL1A1^LOW^ myCAF in epithelium-adjacent regions (**Fig. 3D-E**). At the invasive stage, the stromal distribution was less associated with tumor distance, except for CD105^pos^/COL1A1^MID^ fibroblasts, which were enriched in the distal stroma. We also analyzed immune and vessel composition relative to epithelial distance and observed that vessels were more common distally at LG, but unassociated with distance in HG and INV (**Supplementary Fig. 2**). Other immune cells dominated the peritumoral region at both LG and HG, and myeloid cells were enriched near the epithelium, slightly at LG and strongly at INV.

Given this distance-dependent bias in stromal identity, we tested whether epithelial phenotype influenced the identity of adjacent subepithelial fibroblasts. We created a mask-constrained radius search algorithm to identify every stromal cell within 50 μm of each epithelial cell, enabling the characterization of the average stromal niche surrounding each epithelial subpopulation (**Supplementary Fig. 3A** and **METHODS**). Our results demonstrate that, in addition to shifting with grade, stromal niche composition varied in association with the epithelial subpopulation in proximity (**Supplementary Fig. 3B**). For instance, at high-grade, niches of the CD166^HIGH^/CD133^HIGH^/S100P^HIGH^ and CD166^MID^/CD133^HIGH^ epithelial subpopulations were enriched in CD105^pos^/COL1A1^LOW^ myCAFs compared to those of the other epithelial subpopulations. Immune and vessel composition also varied relative to the neighboring epithelial subpopulation (**Supplementary Fig. 3C**). Altogether, these data suggest that epithelial-stromal distances strongly orchestrate fibroblast identity and significant interactions within the subepithelial regions.

### Xenium *In Situ* spatial transcriptomics of an IPMN progression dataset highlights the key role of epithelial signatures and myCAFs during disease progression

To further investigate fibroblast heterogeneity and the subepithelial changes, we performed Xenium *in situ* imaging on 4 TMAs (32 human pancreas samples, 18 patients) covering normal, ADM, LG, HG and INV (**Fig. 4A**, **Supplementary Table 1**). We used the immune-oncology panel and an additional 100-gene custom panel for epithelial stem and fibroblast lineages to spatially resolve 480 genes at single-cell resolution (see **METHODS** and **Supplementary Table 3**). We performed single-cell segmentation using Proseg and our custom anucleated segmentation pipeline and performed cell typing by unsupervised clustering in two steps, identifying Crude (**Fig. 4B**) and Refined clusters (**Fig. 4C**). We controlled batch effects, and cluster purity by calculating Local Inverse Simpson’s Index (iLISI and cLISI scores, **Supplementary Fig. 4**). Our anucleated segmentation pipeline increased cellular capture, especially fibroblast content (**Supplementary Fig. 5**). Clustering spatial overlay of crude clusters on representative tissue sections confirmed proper cell type assignment (**Fig. 4B**). Refined clustering of the initial crude clusters revealed multiple epithelial, stromal, immune, neuroendocrine, and endothelial subpopulations (**Fig. 4C**). Specifically, it resolved 15 epithelial and 5 stromal subpopulations annotated based on their top differentially expressed genes (**Fig. 4D-E**, **Supplementary Table 4**). One stromal subpopulation, lacking defining markers, was labeled Unclassifiable_Fibroblasts. For the epithelial compartment, we further informed annotations by *in situ* localization of each cluster under the supervision of a board-certified pathologist, distinguishing clusters associated with benign histological features from those located in pre-cancerous or cancerous regions. To identify cell populations associated with disease progression, we fitted a mixed-effects model for each refined cell type (**Fig. 4F**). The strongest positive associations were found for myCAF and iCAF in the stroma, and for Tumoral_CXCL12_REG4, followed by Rare_PTEN_GLUL and Tumoral_Invasive_S100P_CD166_CEACAM6 in the epithelium. Conversely, benign interlobular/intralobular ducts, benign duct-stem, and vascular-associated stromal cells were progressively depleted. This shift mirrored the overall cellular composition across grades: benign ductal and acinar states gave way first to goblet and mucinous tumoral cells, then to invasive tumoral cells (**Fig. 4G**). In the stroma, iCAF emerged at ADM and persisted through INV, while myCAF expanded markedly from LG to INV and the vascular-associated compartment reduced with disease progression (**Fig. 4H**).

**Figure 4.**
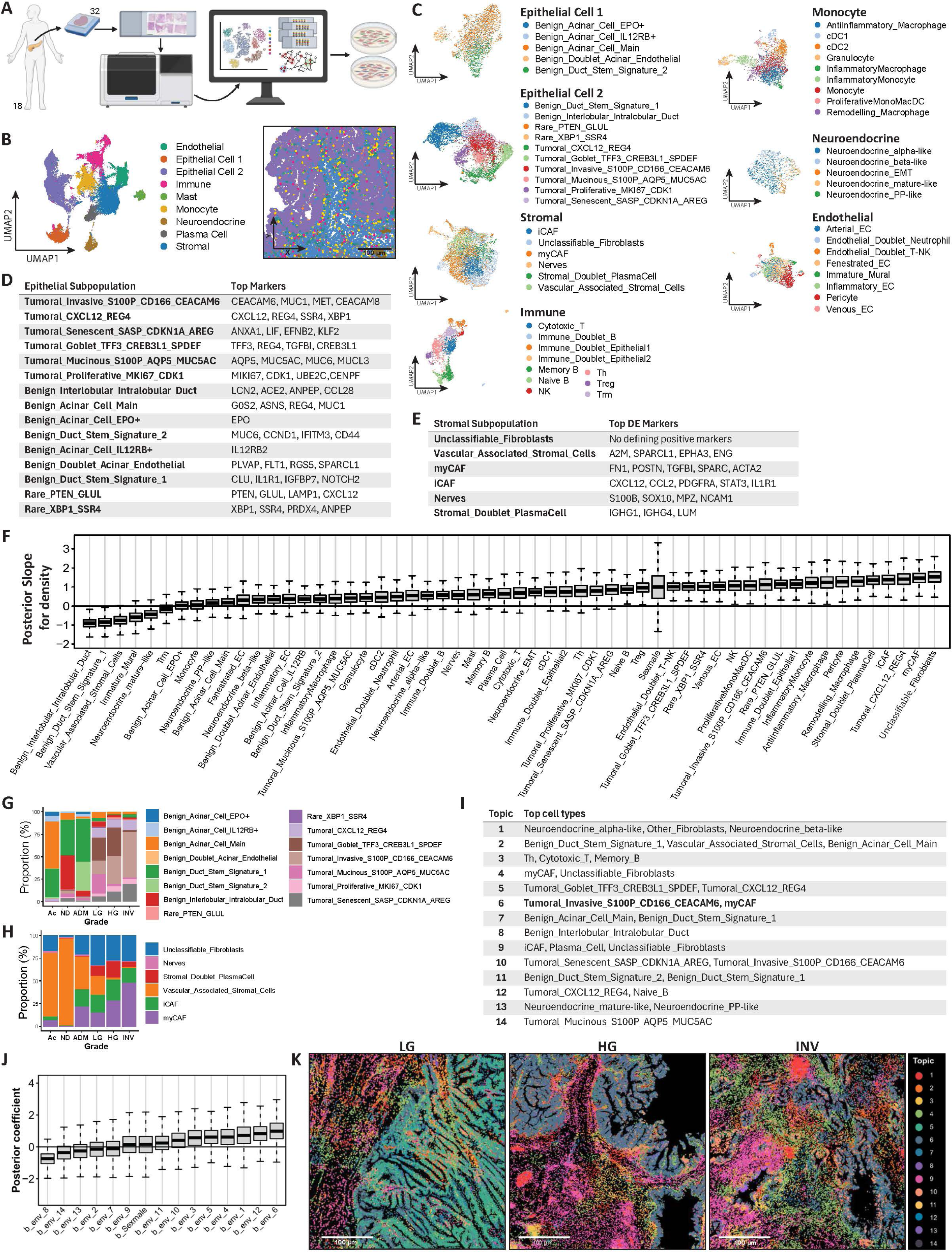
Xenium In Situ spatial transcriptomics of an IPMN progression dataset highlight the key role of epithelial stem cell signatures and mycafs during disease progression. **A.** Schematic overview of the experimental design for Xenium in Situ sample preparation, data extraction and analysis towards *in vitro* evaluation. Created in Biorender. **B.** UMAP projection of single cells, down sampled from n=32 samples, colored according to crude clustering results, annotated per cell type. Associated *in situ* projection of x, y cell positions in HG region, colored according to crude cell type. **C.** Multiple UMAP projections of single cells from initial crude clusters (Epithelial Cell 1, Epithelial Cell 2, Stromal, Immune, Monocyte, Neuroendocrine, and Endothelial), colored according to refined clustering results, annotated by sub cell types. **D.** Table of top DE expressed genes per epithelial refined cluster. **E** Table of top DE expressed genes per stromal refined cluster. **F.** Estimated per-cell-type posterior slopes for Bayesian hierarchical mixed-effects modeling of cell type densities, adjusted for Sex and Sample. G. Stacked bar graph of cell proportion of each epithelial cell refined cluster per grade. **H.** Stacked bar graph of cell proportion of each stromal cell refined cluster per grade. **I.** Table of top cell types (>10% cell proportion) in each Topic from SpaTopic analysis across samples. J. Estimated per-topic posterior slopes for Bayesian hierarchical ordinal regression of topic densities, with every topic as an independent fixed effect, adjusted for sex and sample. **K**. Representative LG, HG, and INV tissue sections showing localization of cells colored by Topics type. LG - low-grade dysplasia, HG - High-grade dysplasia, INV - invasive cancer.

To map regional patterns, we analyzed samples using SpaTopic. We identified 14 global cellular patterns enriched in different proportions of each subpopulation (**Fig. 4I** and **Supplementary Fig. 6**). Modeling topic association with disease progression revealed Topic 6, composed mostly of Tumoral_Invasive_S100P_CD166_CEACAM6 and myCAFs, as the most associated (**Fig. 4J**). Spatial overlay on representative sections confirmed Topic 6 enrichment from LG to HG-INV (**Fig. 4K**). To test tumor-stromal interactions directly, we performed co-localization and ligand-receptor analyses. Pairwise co-localization was dominated by homotypic aggregation and lineage-restricted neighborhoods and did not resolve a specific tumor-stromal pairing at any grade (**Supplementary Fig. 7A-D**). Ligand-receptor inference focused on Topic 6 and then resolved an epithelial-fibroblast interface within an otherwise dense communication network. Tumor cells engaged mediators such as FN1, EFNB2, TGFB1, and TGFBI in fibroblasts (**Supplementary Fig. 8** and **Supplementary Table 5**). Together, these data highlight the central role of myCAF in disease progression, in coordination with adjacent epithelial components.

### Subepithelial fibroblasts co-expressing NOTCH1 and NESTIN are a prevalent subepithelial population associated with tumor progression

To resolve the molecular programs engaged in the subepithelial stroma during tumor progression, we analyzed distance-resolved gene expression in the Xenium dataset. We applied our mask-constrained nearest neighbor algorithm (**Fig. 3A**) and found associations between stromal phenotype distribution and distance to the nearest epithelial cell, replicating our CyCIF observations (**Fig. 5A**). The myCAF proportion was the highest in tumor proximal regions and became the most prominent population at the tumor edge from LG to HG. We analyzed single-gene expression in stromal cells across distance bins from the epithelium and found 208 genes peaking in the 0-50 μm range across all grades, while 72 peaked at 500-1000 μm (**Fig. 5B**). By comparing the 0-50 μm bin against distal bins, we identified 105 differentially expressed genes (DEGs) shared by LG and HG. Among the shared genes, a subset showed greater differential expression in HG than in LG, spanning stromal progenitor (*NOTCH1*, *NES*), mesenchymal transcription factor (*SOX4*, *ZEB1*), cell cycle (*CCND1*, *CDK6*, *RB1*), interferon-responsive (*STAT1*, *IRF2*, *IL1R1*), and antigen-presenting-like (*CD4*, *CD74*) programs (top 25 genes in **Fig. 5C**).

**Figure 5.**
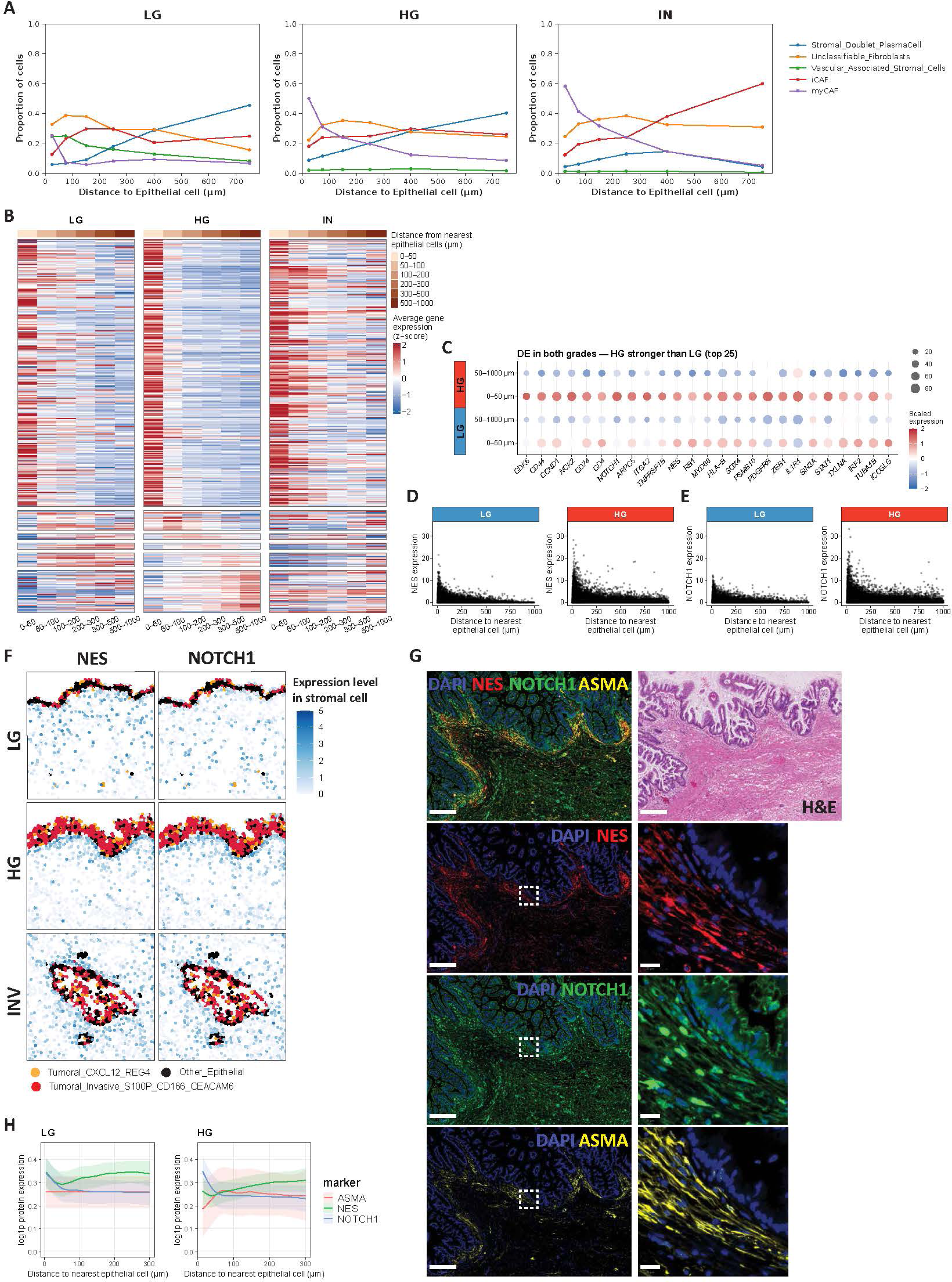
Subepithelial fibroblasts co-expressing NOTCH1 and NESTIN are a prevalent subepithelial population associated with tumor progression. **A.** Connected dot plot of proportions of stromal refined clusters as a function of stromal cell distance to nearest epithelial cell, calculated with our nearest neighbor algorithm. **B.** Heatmap of mean gene expression per distance to nearest epithelial cell bin (0-50, 50-100, 100-200, 200-300, 300-500, and 500-1000 μm, expression z-scored per gene across all grades, total of 348 genes detected in at least 5% of cells). **C.** Dot plot of top 25 genes significantly upregulated in stromal cells within 0-50 μm tumor edge (relative to other distance bins) in both LG and HG, and showing stronger edge versus rest Log2 fold change in HG than LG (for genes detected in at least 40% of cells). **D.** Dot plot of NES expression in stromal cells as a function of distance to nearest epithelial cell (in μm), for both LG and HG. **E.** Dot plot of NOTCH1 expression in stromal cells as a function of distance to nearest epithelial cell (in μm), for both LG and HG. **F.** Representative LG, HG, and INV tissue sections showing localization of epithelial cells (colored by refined cluster: Tumoral_Invasive_S100P_CD166_CEACAM6 cells in red, Tumoral_CXCL12_REG4 cells in orange, and others in black), and stromal cells only, colored by expression levels of NES or NOTCH. G. Representative HG tissue section showing immunofluorescence signal for DAPI (blue), NES (red), NOTCH1 (green), and ASMA (yellow) (scale bar 250µm), with zoomed views (scale bar 20µm), and H&E-stained subsequent tissue section. **H.** Generalized additive model of protein expression (Log1p) relative to distance to nearest epithelial cell, respectively for LG and HG ROIs. Lines are across-ROI average trends with 95% confidence interval of the mean. LG - low-grade dysplasia, HG - High-grade dysplasia, INV - invasive cancer.

We focused on *NOTCH1* and *NES,* an intermediate filament protein marker of progenitor cells, given our limited panel and as previous studies have shown that the NOTCH intracellular domain directly activates *NES* expression in gliomas (30). *NES* and *NOTCH1* transcripts peaked in stromal cells immediately adjacent to the epithelium and declined with distance, with a sharper proximal peak at HG than LG (**Fig. 5D–E**). Spatial overlays confirmed this subepithelial enrichment of *NES* and *NOTCH1*, which intensified from LG to HG and remained tumor-proximal at the INV stage (**Fig. 5F**). We next validated these observations at the protein level by immunofluorescence, which identified subepithelial fibroblasts co-expressing NES, NOTCH1, and αSMA (**Fig. 5G**). Immunofluorescence quantification by distance recapitulated the transcriptomic gradients (**Fig. 5H**). NOTCH1 protein expression decreased with distance from the epithelium at both LG and HG, while NES was highest in the immediate subepithelial stroma before declining and increasing again farther out. These findings define a subepithelial NOTCH1-NES program in subepithelial fibroblasts that intensifies with grade.

### Nestin-expressing fibroblasts emerge in high proximity to tumor cells *in vitro* and Notch1 inhibition modulates spatial tumor growth patterns in co-cultures

To characterize the epithelium-proximal cells expressing high levels of *NES* and *NOTCH1*, we investigated *NES* and *NOTCH1* expression levels in each subpopulation. Across most stromal subpopulations, *NES* and *NOTCH1* expression increased toward the tumor, but the magnitude of this distance dependence was subpopulation-specific. MyCAF displayed among the steepest proximal gradients for both genes (**Fig. 6A**). Gating both markers within the 0-100 μm range defined a new population termed sub-epithelial NES/NOTCH1 fibroblasts (**Fig. 6B-C**), 67.5% of which had been previously annotated as myCAF, suggesting the NES/NOTCH1 population overlaps subsequentially with the myCAF phenotype (**Fig. 6D**). Sub-epithelial NES/NOTCH1 fibroblasts are enriched from LG to HG, and less prominent in INV, while myCAF are enriched throughout progression, at both 0-50 and 50-100 μm ranges (**Fig. 6E**). Beyond *NES* and *NOTCH1*, this population was distinguished from other stromal cells by a broader signature including *RGS5, FLT1, PLVAP, EFNB2, ICOSLG, ITGA1/2, NOTCH3, TGFB1,* and *PDGFRB* (**Fig. 6F**). Gene set enrichment analysis associated this population with proliferation regulation, regulation of cell differentiation, and immune system processes (**Fig. 6G**).

**Figure 6.**
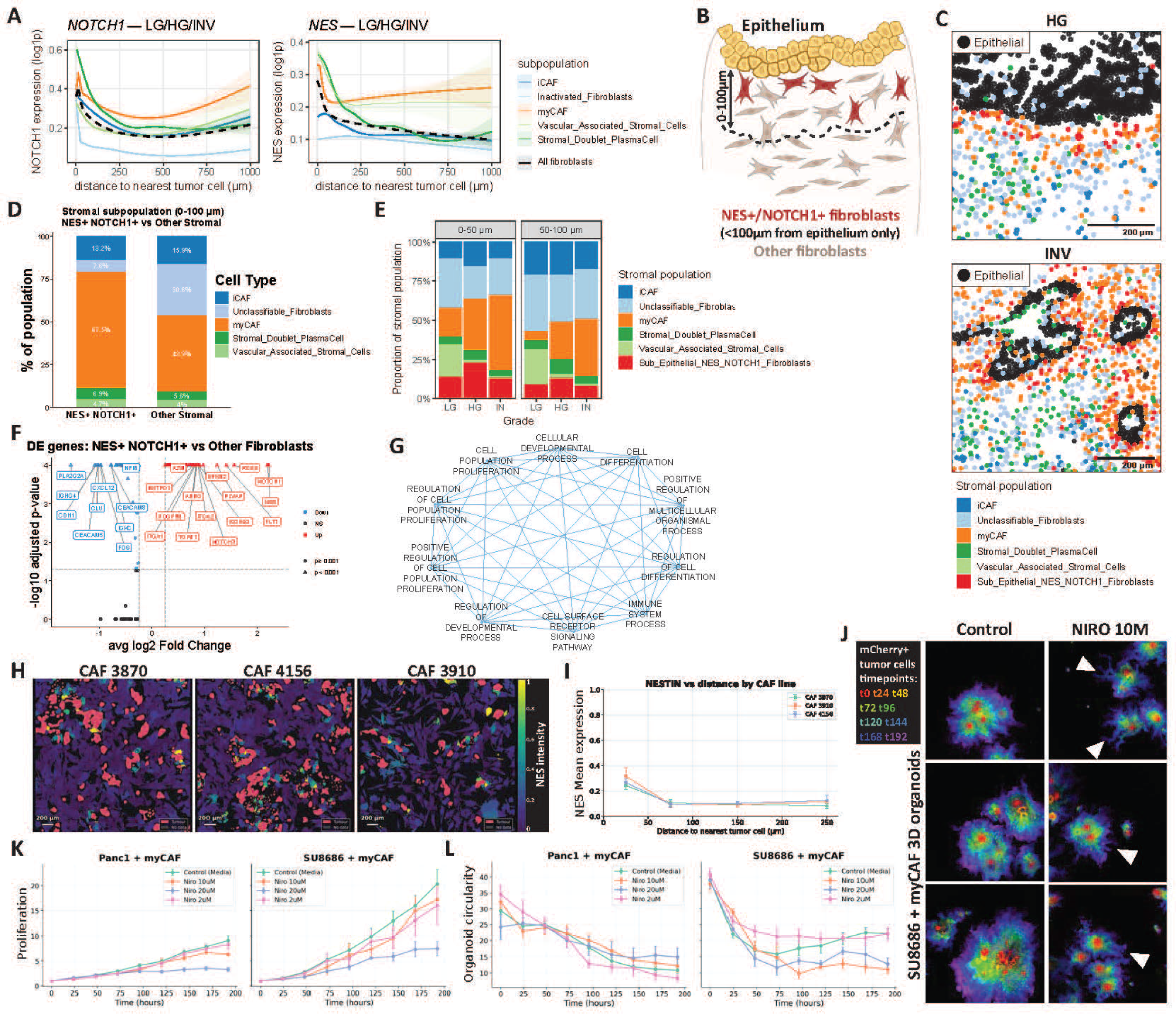
Nestin-expressing fibroblasts emerge in high proximity to tumor cells *in vitro* and Notch1 inhibition modulates spatial tumor growth patterns in co-cultures. **A.** Generalized additive model estimations of smoothed conditional mean of stromal NES and NOTCH1 expression levels (log1p) relative to distance to the nearest epithelial cell, per stromal population (colored lines), or for all populations (black dashed line). Shaded areas are 95% confidence intervals (CI) for conditional mean. **B**. Schematic overview of NES+/NOTCH1+ fibroblasts within 0-100 microns from the epithelium. Created in Biorender. **C**. Representative HG and INV tissue sections showing localization of epithelial cells in black, and stromal cells only, colored by refined clusters including subepithelial NES^HIGH^/NOTCH1^HIGH^ population. **D**. Bar plot of initial refined population proportion per NES^HIGH^/NOTCH1^HIGH^ population or other populations. **E**. Stacked bar plot of stromal population proportion per grade within 0-50 or 50-100 μm from epithelium. **F**. Volcano plot of deferentially expressed genes in subepithelial NES^HIGH^/NOTCH1^HIGH^ population versus other populations. **G**. GO term enrichment network for top genes significantly upregulated in subepithelial NES^HIGH^/NOTCH1^HIGH^ population versus other populations (Log2 fold change > 0.25). **H**. Representative images of 2D co-cultures of SU8686 and CAF lines showing NES intensity in CAF and tumor cells in pink, for respectively CAF 3870, 4156, or 3910. **I**. Connected dot plot of NES expression level according to distance to nearest tumor cell in 2D co-cultures of SU8686 and CAF lines (CAF 3870, 4156, or 3910). **J**. Composite images of mCherry+ tumor cells at different timepoints, from SU8686 + myCAF 3D organoids, untreated controls and Nirogacestat 10M treated. Composite images are overlays of all timepoints with the earliest in the foreground (T0-T192). **K**. Connected dot plot of SU8686 proliferation according to time, from SU8686 + myCAF 3D organoids with untreated control, or Nirogacestat treatment (2, 10, and 20M). **L**. Connected dot plot of organoid circularity according to time, from SU8686 + myCAF 3D organoids with untreated control, or Nirogacestat treatment (2, 10, and 20M). LG - low-grade dysplasia, HG - High-grade dysplasia, INV - invasive cancer.

To test whether tumor cell proximity induces NES expression we co-cultured mCherry-labeled SU8686 pancreatic cancer cells with primary human CAF (lines 3870, 4156, and 3910) and performed immunofluorescence for NES. We observed more NES expression in CAF in proximity (< 50 µm) to tumor cells, for each of the CAF lines tested (**Fig. 6H-I**). To functionally assess whether Notch signaling supports tumor growth, we co-cultured mCherry-labeled PANC1 or SU8686 pancreatic cancer cells with primary human myCAF (line 3870) and treated them with the γ-secretase/Notch inhibitor nirogacestat. Notch inhibition reduced co-culture expansion in a dose-dependent manner, with the strongest effect at 20 µM for both tumor lines (**Fig. 6J-K**). Colony circularity declined as colonies spread over time (**Fig. 6L**). In SU8686, 10-20 µM nirogacestat further reduced circularity from 100h onward, with maximal effect by 200h. On representative images overlaying mCherry+ tumor cell signal at each time point, Notch inhibition yielded smaller, less cohesive colonies with prominent elongated projections (**Fig. 6J**). These co-culture results indicate that Notch signaling supports tumor growth in tumor cells co-cultured with myCAFs.

## DISCUSSION

The role of the tumor microenvironment and particularly CAFs in PDAC is increasingly established (17–19). Nonetheless, little work has assessed fibroblast heterogeneity and its role in IPMN progression and IPMN-derived PDAC. In this study, we combined methods in computational pathology, multiplex imaging, and spatial transcriptomics to identify key fibroblast populations that progress during disease.

We first leveraged a method that combines classical computational pathology with a typical bioinformatic pipeline used in single-cell biology. This allowed us to screen 343 whole-slide images from 83 patients, leveraging epithelial and stromal heterogeneity, which could then be further interrogated at the protein and RNA level using spatial biology methods. All three approaches highlighted similar progressive patterns with disease progression: epithelial and stromal remodeling. Stromal findings have been previously described, but mostly restricted to immune populations (8, 31, 32). They described enhanced immune surveillance observed in LG compared to HG IPMN, and progression towards a more immunosuppressive TME with macrophages and T cells alterations.

Interestingly, from our Xenium data, populations most associated with progression were not epithelial nor immune, but rather the fibroblastic compartment. Studies on fibroblast spatial diversity in pancreatic precursors mostly investigate PanINs. Bell *et al.* demonstrated that fibroblasts harboring features close to PDAC-CAFs are located close to PanINs, suggesting early modulation of TME composition (33). On the other hand, Elhossiny *et al*. described an asynchronous transition of epithelium and stroma during tumorigenesis (34). They observed a contrast between the epithelial evolutionary trajectory and TME composition, with differences between PanIN, resembling more normal ducts, and invasive TME having specific fibroblast populations. Our IPMN data depict a progressive remodeling of both epithelial and stromal compartments with progression while highlighting the importance of distance.

We leveraged clinical and biological advantages of IPMNs in our study. Their epithelial lining is organized in a more linear fashion against the stromal wall, from which emerges a stromal gradient that is thicker than PanINs. Because the IPMN cyst wall is thick, it offered a tractable geometry in which to measure gradients extending from the epithelium into the stroma. This allowed us to study fibroblast diversity in relation to the epithelial component, notably at the pre-invasive stage. Indeed, our spatial statistics methods suggest that the fibroblast gradients emanating from the epithelial components are mostly maintained in the pre-invasive stage.

The existing principle for spatial CAF subtype contextualization in PDAC was set by Ohlund *et al.,* describing myCAF closer to the tumor and iCAF further away (25). Our study is more extensive in terms of the protein and transcriptomic marker panel created to delineate the diversity of CAF. Yet, the dominance of myCAF near the tumor and iCAF further away is replicated both in our proteomic and transcriptomic data. Peng *et al*. had explored peritumoral regions of HG-IPMN and invasive PDAC and showed greater abundance of myCAF than iCAF in tumor edges at both grades (35). While other studies focused only on adjacent stroma, without comparison to stromal cells located further away from the tumor, we evidence the existence of a gradient effect driving fibroblast phenotype as early as LG-IPMN, and strongly in HG. Gradient effects are preserved, but diminished, in invasive disease, which is likely due to the more aggressive and disseminative nature of PDAC.

Our spatial statistics-guided methods highlight a critical area of disruption right at the junction between epithelial cells and the stroma. Multiple genes defined this population; however, *NES* and *NOTCH1* stood out as key markers that identify fibroblasts expanding during low- to high-grade transition. One study by Nielsen *et al*. observed higher NES expression in juxtatumoral CAFs compared to peripheral CAFs in human PDAC (36). Our work is the first description of subepithelial NES/NOTCH1 CAF establishment in IPMNs, prior to the invasive stage. NES and NOTCH1 expression were observed in multiple other organs or pathologies (37, 38) and it was shown that the NOTCH intracellular domain activates directly *NES* expression in gliomas (30). Concerning NOTCH1, most of the focus was on its role in cancer cells; however, recent studies evidence its role in the TME (39). In lung cancer, NOTCH1 silencing in CAF reversed the antiproliferative and proapoptotic functions of the conditioned media of CAF exposed to apoptotic cancer cells (40). In CAF, intracellular NOTCH1 dictates stemness and plasticity of melanoma initiating cells and regulates melanoma aggressiveness (41). Increased NOTCH signaling in CAF reduced melanoma growth, thus conferring tumor-suppressive functions (42). NES is upregulated in lung ASMA+ myofibroblasts and plays a pro-fibrotic role by facilitating TGFβ type I receptor (TβRI) recycling through Rab11 (43). In liver fibrosis, NES is induced in hepatic stellate cells by TGFβ, and knocking down NES resulted in caveolin 1-mediated TβRI degradation, reducing TGFβ-induced fibrosis. Although not tested in a cancer context, those studies link NES and fibrosis capacity, known for participating in both pro-tumoral and tumor-restraining functions (44). Therefore, the NES/NOTCH1 axis could be a novel targetable fibrosis-orchestrating event.

This fibroblast population is defined not only by NES and NOTCH1 expression, but also by proximity. This is not surprising given the juxtracrine aspect of contact-dependent NOTCH signaling (39). We could induce this spatially-bound interaction by co-culturing tumor cells with primary myCAFs. A co-submission with Ibanez-Rios *et al*. characterizes the mechanisms underlying the induction of the NESTIN-expressing phenotype in fibroblasts and the key role of growth factors in inducing and NOTCH1 in maintaining this fibroblast state. However, consistent with the general findings of this study, this is likely a transient state owing to the plastic nature of tumor stroma. Our findings suggest that, across fibroblast programs, distance to epithelial cells remains critical in determining identity.

There are limitations to our studies. Spatial biology methods applied to human samples offer unprecedented clinically relevant and native spatial architecture but are costly and limited the number of samples we could analyze. Computational pathology allowed us to consider a larger cohort, observe general spatial organization changes, and then investigate them at transcriptomic and proteomic levels. We established that fibroblast identity is organized relative to distance from the neoplastic epithelium using three approaches and validated it in an independent cohort and *in vitro* to compensate for sample size. Another limitation is that human tissues provide only a static snapshot, so progression cannot be observed longitudinally; therefore, grade-specific associations could only be inferred from cross-sectional comparisons. We obtained the CyCIF and Xenium datasets from a limited target panel. Further validation using genome-wide spatial methods, when available, would strengthen our understanding. To further study signatures associated with the sub-epithelial NES/NOTCH1 fibroblasts, we performed artificial gating of the sub-population, which was not natural. The reality is that CAF phenotypes are much more complex due to their extreme context-dependent plasticity.

Overall, our data shows pre-invasive molecular reprogramming with microenvironment alterations prior to invasive transition. Findings suggest extreme plasticity of the stroma and the capacity of tumor cells to reorganize their immediate microenvironment. Our findings indeed further highlight novel therapeutic targets that could control disease and improve current therapeutic approaches.

## METHODS

### Cell-centered clustering computational pathology processing pipeline

Whole-slide images (WSIs) generated during routine clinical care were collected from the Centre Hospitalier de l’Université de Montréal digital pathology image repository. Region of Interest selection was performed by two board-certified pathologists unaware of the research aims and asked to randomly select representative areas approximately 500 x 500 µm in size on the outer edge of the cystic process in QuPath 0.5.1 (45). These regions were positioned so that the epithelial lining would be oriented to one side of the annotation and a consistent wall of stroma was bound to this lining. Regions of interest were exported as ome-tiffs.

We implemented a four-stage, cell-centered representation-learning pipeline on the exported ome-tiffs (**Fig. 1A**): (1) nuclear segmentation of whole-slide H&E images; (2) patch extraction centered on each detected nuclear centroid; (3) embedding of every tile with the UNI2-h pathology foundation model; and (4) unsupervised k-means++ clustering of the resulting feature space with UMAP-based visualization. All slides were pooled prior to clustering so that cluster identities are shared across the cohort. Nuclei were detected in QuPath v0.6 using the StarDist2D extension and the pre-trained he_heavy_augment model (45, 46). Prior to inference, stain intensity was normalized per image only using percentile-based scaling (0.2nd and 99.8th percentiles) computed on a downsampled representation of the whole image (maximum dimension 4096 pixels).

Rather than using a regular grid, we extracted a single 224 x 224-pixel patch in full-resolution JPEG format centered on each detected nuclear centroid at a 40X magnification (0.23 µm/pixel) along with corresponding (x,y) slide-level coordinates. Nuclei close to slide image boundaries resulting in incomplete patches were excluded. Each cell-centered patch was embedded using UNI2-h (4), a 1,536-dimensional vision transformer pretrained by self-supervised learning on histopathology images using 224 x 224-pixel patches natively and relying on DINOv2 with self-distillation with multi-crop augmentation, masked image modeling, and KoLeo regularization (47, 48). Weights were obtained from the official UNI2-h model repository and loaded using timm library according to the authors’ published configuration. No stain normalization or color augmentation was applied. Inference was performed in evaluation mode without gradient computation, using automatic mixed precision (float16) on a single NVIDIA H100 80 GB GPU with PyTorch implementation (49). The 1536-dimensional class-token output was taken as the cell-centered patch representation. Embeddings were written to HDF5 alongside a CSV export retaining the tile path for each row. All patch-level embeddings from all slides were pooled into a single matrix of shape 777,710 x 1536, allowing shared cluster representation across all regions of interest.

Patch-level clustering was performed in Python using the cuML GPU implementation of the scalable k-means++ method, through a range of k=5 to 100 in increments of 5, selecting the elbow of the inertia curve located as the point of maximum perpendicular distance from the chord joining the first and last points of the min-max-normalized curve (50). A fixed seed of 42 was used for all downstream analyses. The final partition was obtained by fitting k-means++ at k = 35, determined as above, with a maximum of 300 iterations. Other internal indices (silhouette, Calinski-Harabasz, Davies-Bouldin) did not identify a single optimum through the range of k=5 to 100.

Clustering was used as an exploratory and descriptive device: no downstream statistical claim depends on the specific value of k. For visualization only, the 1536-dimensional embedding patch representations were projected to 2D UMAP with 30 nearest neighbors, a minimum distance of 0.1, set with Euclidean metric, alongside Matplotlib and Glasbey palette.

To review cluster patch histopathological morphology without biasing inspection toward dense regions of the feature space, we sampled 1000 patches per cluster using a coverage-based rather than a random strategy. Within each cluster, k-means++ implemented as above with k = 1000, with a maximum of 300 iterations, was applied to the two-dimensional UMAP coordinates of that cluster’s members, and the tile nearest each resulting centroid (by squared Euclidean distance) was selected, with duplicate assignments resolved by taking the next-nearest unselected tile. This method allows more substantial sampling of sparse and peripheral regions of each cluster than uniform random sampling would, while remaining denser towards each cluster’s core. Following full sampled patch joint review by cluster by two board-certified pathologists (A.K. and V.Q.T.), each cluster was assigned a consensus morphological semantic description and a corresponding semantic annotation of grade-specific atypia or benign cluster state and further classified as predominantly epithelial versus stromal.

### CyCIF of IPMNs and PDAC

For CyCIF, we collected samples from routine surgeries at Vanderbilt University Medical Center. These samples were fixed in formalin and embedded in paraffin, then sectioned using a microtome (4 microns). They came from 16 patients with IPMN or IPMN associated with PDAC and covered the spectrum of pancreatic tumorigenesis: normal pancreatic tissue (N), acinar-to-ductal metaplasia (ADM), low-grade IPMN (LG), high-grade IPMN (HG), and PDAC (INV). Clinical data is available in **Supplemental Table 1**.

Whole slide image (WSI) sections from 16 formalin-fixed paraffin-embedded (FFPE) samples were stained with a validated 19-plex marker panel (see **Supplemental Table 2**) and scanned with Axio Scan.Z1. The CyCIF protocol includes tissue deparaffination with xylene washes (3 x 3 min), and rehydration with ethanol (1 x 2 min 100% EtOH, 1 x 2 min 95% EtOH, 1 x 2 min 70% EtOH), and doubly distilled water (ddH2O, 1 x 2 min). For heat antigen recovery, the slides were placed in a polypropylene Coplin jar immersed in 10 mM Tris buffer, 1 mM EDTA, pH 9, and heated in a 95°C water bath for 5 minutes, then cooled for 30 min, and washed in PBS for 2 x 3 min. Tissues were blocked for 10 minutes with Superblock (#NC9782835, Scytek Laboratories). Primary antibody combinations with distinct fluorophores are diluted in Agilent antibody diluent (#S0809, Dako). 100 µL of diluted antibody is applied to each slide for overnight incubation at 4°C in a humid chamber. Slides are washed 3 x 3 min in PBS, then stained with DAPI (#D9542, Sigma-Aldrich) for 10 min, and mounted in 50% glycerol (#56-81-5, BioShop) in PBS. Slides are imaged with the Axio Scan.Z1 (Zeiss, Oberkochen, Germany) for DAPI, 488 nm, 555–594 nm, and 647 nm channels. The coverslips are removed from the slides and washed 3 x 3 min in PBS to remove residual glycerol. The slides are incubated for 2 hours in 3% hydrogen peroxide, pH 9, between two high-power LED lamps (B08D37RNXM, MEIKEE International, Hong Kong, China), each generating 15,000 lumens at the top and bottom of the slides. The slides are examined by DAPI at 488 nm, 555 nm, and 594 nm to detect any residual staining. We then proceed with blocking, primary antibody incubation, scanning, and fluorescent signal removal for the following antibody combinations.

### CyCIF data processing

Quality control of raw images for shading correction was performed with ZEN 3.8. We used the open-source image processing software Qupath (version 0.5.1) (45). We performed WSI-level image registration using the ImageCombiner-0.3.0 extension. We annotated regions of interest (ROIs) of about 1×1 mm, in each sample (see **Supplemental Table 2**), representing tissue structures, including the adjacent stroma, from a single stage of the disease (normal duct, acinar-to-ductal metaplasia, low-grade dysplasia, high-grade dysplasia, or invasive cancer). Annotation was supervised by a board-certified pathologist (Dr. Vincent Quoc-Huy Trinh). ROIs were exported as .TIFF, and a second image registration was performed using Warpy interactive image combiner to correct minimal shifts between image channels from the previous WSI registration. Cell segmentation was performed using Qupath (45), extracting the mean and standard deviation of pixel intensities of each subchannel (marker) within each cellular segment, and x,y cell centroids. Spatially resolved single-cell data was exported as .csv.

### CyCIF cell phenotyping and analysis

Under the supervision of a board-certified pathologist, tissue structures corresponding to epithelial or tumor cells were manually annotated to distinguish epithelial and tumor cells from stromal cells. This is done by recognizing histological characteristics and localizing marker expressions. For each image, the annotation is exported as a binary mask. We normalized immune and vascular markers (CD45, CD3, CD4, CD8, CD11b, CD163, CD20, and CD31) by dichotomizing signals using fluorescence intensity thresholds for positivity. For other markers (S100P, SPP1, CD44v9, CD166, CD133, CD105, αSMA, CD74, COL1A1, CXCL12, and FAP), we kept continuous expression values and normalized signals by measuring the background fluorescence signal and subtracting it from the mean intensity of each cell, independently for each image. We limited the outlier signal to the [Quartile 3 + 1.5x(Inter Quartile Range)] and rescaled it to the [0,1] range using the MIN-MAX values for each marker. Tissue autofluorescence may generate false-positive signals; we detected them using a logical conjunction. If the cell intensity exceeded the established threshold for all markers during the first three staining cycles, we classified the cell as non-specific. Thresholds were established for the markers CD4, CD45, CD11b, CD166, and CD3, based on the visual identification of erythrocytes and non-specific fluorescence. For phenotyping single cells, we implemented our R pipeline that considers marker expression levels and cell presence in the annotated tumor mask. Cells classified as non-specific (autofluorescence) are eliminated unless they are CD31+, in which case they are assigned to the endothelial type. CD45+ cells are classified as immune cells. CD3 is used to identify lymphocytes. CD45+/CD3+/CD4+/CD8+ cells are assigned to double-positive T lymphocytes (DP T lymphocytes). CD45+/CD3+/CD4+ cells are assigned to helper T lymphocytes. CD45+/CD3+/CD8+ cells are assigned to cytotoxic T lymphocytes. CD45+/CD3+/CD4-/CD8- cells are double-negative T lymphocytes (DN T lymphocytes). CD20 is used for the identification of B lymphocytes; CD45+/CD20+ cells are assigned as B lymphocytes. CD11b characterizes the myeloid lineage: CD45+/CD11b+/CD163+ are assigned to the M2 macrophage, and CD45+/CD11b+ to the myeloid cell type. For CD45^neg^ - non-immune cells, if the cell’s x,y position falls within the annotated tumor mask, we assign it to the epithelial/tumor cell type. Otherwise, CD31+ cells are typed endothelial, and CD31^neg^ cells are non-immune/non-endothelial stromal cells, considered fibroblasts in our study. A second step for subpopulation typing of epithelial and stromal cells is performed independently using Leiden unsupervised clustering from Scanpy in Python, considering epithelial markers CD133, CD166, SPP1, CD44v9, and S100P, and stromal markers CD105, αSMA, CXCL12, FAP, CD74, CD105, and COL1A1, respectively. UMAPs, matrix plots, and other data visualizations were done using scanpy, matplotlib, ggplot2.

### Distance-based algorithms

We developed two algorithms to specifically investigate stroma as a function of adjacent epithelial cells using distance-based approaches. The first algorithm is a Mask-Constrained Nearest Neighbor Algorithm. The goal is to assign each fibroblast its distance to the nearest epithelial/tumor cell to study the fibroblast phenotype as a function of its distance from the epithelium. The algorithm relies on the brute-force calculation of the Euclidean distance between each cell in the stromal mask and all cells in the epithelial/tumor mask. For each stromal cell, the smallest distance is identified (minimum search principle), corresponding to the distance to the nearest epithelial/tumor neighbor. The computation is vectorized on a GPU (using CuPy), and distance calculations are performed in parallel to accelerate processing. The second algorithm is a Mask-Constrained Radius Search Algorithm. The goal is to assign to each epithelial cell the fibroblasts whose distance to that same cell does not exceed a fixed maximum distance. This allows us to study the fibroblast phenotype based on the nearby epithelial type. The algorithm uses brute-force range finding, adding a new constraint for each attribute (belonging to the epithelial/tumor or stromal mask), and calculates Euclidean distances. The range finding is performed only from the epithelial mask to the stromal mask. The computation is vectorized on a GPU (using CuPy), and distance calculations are performed in parallel to accelerate processing. Both algorithms were implemented in Python and use scanpy, pandas, and numpy libraries. Algorithms were adapted for Cell-centered clustering computational pathology processing pipeline output, CyCIF data, Xenium data, and *in vitro* co-cultures immunofluorescence data.

### Xenium in situ imaging

We collected samples form routine surgeries performed at the Centre Hospitalier de l’Université de Montréal. Sections of 4 TMAs of 8 FFPE samples each were mounted on Xenium slides and processed by 10X Xenium *In situ*, according to manufacturer protocol, using the v1 immuno-oncology human panel and custom gene panel of a 100 added genes (clinical data **Supplemental Table 1**, for panel see **Supplemental Table 3**). We annotated regions of interest (ROIs) in each sample, representing tissue structures, including the adjacent stroma, from a single stage of the disease (normal duct, acinar cells, adipose tissue, acinar-to-ductal metaplasia, low-grade dysplasia, high-grade dysplasia, or invasive cancer).

### Xenium cell segmentation pipeline enriched for fibroblasts

We used Proseg and our custom anucleated cell finder for unsupervised segmentation of transcripts. The anucleated Nuclei Identification Pipeline identifies cells whose nuclei were not captured in the imaging plane, but whose cell bodies (and thus, transcript clouds) are still present. This pipeline identifies these cells by finding high-density “transcript shadows” in regions previously classified as background. It takes the outputs from a standard Xenium Ranger and proseg run, identifies new cell candidates, and generates a new transcript file. This new file can then be used as input for a secondary proseg run to fully segment these previously missed cells.

### Xenium cell typing

Segmented single-cell data were processed using Seurat (version 5.3.1) together with custom pipelines for cell clustering and annotation. For each sample, high-resolution clustering was performed, and 1,250 cells were randomly subsampled from each resulting cluster to construct a balanced dataset. This downsampled dataset was then clustered in two stages: an initial low-resolution clustering to identify Crude clusters and a second high-resolution clustering of each Crude cluster to identify Refined clusters. Cluster assignments (both Crude and Refined) were ultimately propagated to the full, non-downsampled dataset using a custom seedCluster() function, based on Euclidian distance. Cell type annotation was based on differentially expressed genes identified with FindAllMarkers() using ROC analysis, for grade-specific annotation of epithelial cells, on *in situ* spatial projection. Cell clusters showing evidence of cross-contamination between distinct cell types were annotated as Doublet.

### Xenium data analysis

Data analyses and visualizations were performed using Seurat, ggplot2, ggven. Mixed-effect modeling was performed with a custom pipeline using brms, posterior, projpred, dplyr, tidyr, and janitor packages. We performed Bayesian hierarchical mixed-effects modeling of cell type densities (Grade ∼ Sex + Density_z + (1|Sample) + (1 + Density_z | CellType)). We treated grades as ordinal (link=“logit”, for N-LG-HG-INV), and the model allowed cell-specific slopes. We estimated per-cell-type posterior slopes from the model and plotted them in order of posterior median. Spatial topic analysis was done using the package SpaTopic, using parameters: ntopics = 14, sigma = 50, region_radius = 100. We performed Bayesian hierarchical ordinal regression of topic densities, with every topic as an independent fixed effect, adjusted for sex and sample. We estimated per-topic posterior slopes from the model and plotted them in order of posterior median. We performed generalized additive model (GAM) estimations of smoothed conditional mean of stromal NES and NOTCH1 expression levels (log1p) relative to distance to the nearest epithelial cell using geom_smooth(). Differential expression genes analysis was done using FindMarkers() and a Wilcoxon test. GSEA analysis was done with G:Profiler and Cytoscape.

### Immunofluorescence of IPMNs and PDAC

Whole slide image (WSI) sections from 38 formalin-fixed paraffin-embedded (FFPE) samples were stained with a validated 3 antibodies panel and scanned with the SLIDEVIEW VS200 (Hamburg, Germany). The IF protocol includes tissue deparaffination with xylene washes (3 x 3 min), and rehydration with ethanol (1 x 3 min 100% EtOH, 1 x 2 min 95% EtOH, 1 x 2 min 70% EtOH), and doubly distilled water (ddH2O, 1 x 2 min). The slides are incubated for 48h in 3% hydrogen peroxide, pH 9, between two high-power LED lamps (B08D37RNXM, MEIKEE International, Hong Kong, China), each generating 15,000 lumens at the top and bottom of the slides for autofluorescence quenching. For heat antigen recovery, the slides were placed in a polypropylene Coplin jar immersed in 50% Sodium citrate pH6 10mM + 50% tris EDTA 1Mm pH9 and heated at 90°C in microwave for 15 minutes, then cooled to room temperature, and washed in PBS for 2 x 3 min. Tissues were blocked for 20 minutes with Superblock (#NC9782835, Scytek Laboratories). Primary antibody combinations [Alexa Fluor 594 anti-Nestin antibody (clone 10C2; BioLegend, cat# 656804, dilution 1:100), Alexa Fluor 488 Alpha-Smooth Muscle Actin Monoclonal Antibody (clone 1A4; eBioscience, AB_2574461, dilution 1:150), Alexa Fluor 647 Notch1 Antibody (Santa Cruz Biotechnology, sc-376403, dilution 1:100)] with distinct fluorophores are diluted in Agilent antibody diluent (#S0809, Dako). 100 µL of diluted antibody is applied to each slide for overnight incubation at 4°C in a humid chamber. Slides are washed 3 x 3 min in PBS, then stained with DAPI (#D9542, Sigma-Aldrich) for 10 min, and mounted in 70% glycerol (#G33-1, Fisher) in PBS. Slides are imaged with the SLIDEVIEW VS200 (Hamburg, Germany) for DAPI, 488 nm, 555–594 nm, and 647 nm channels. Images were annotated in Qupath under the supervision of a board-certified pathologist to create grade-specific ROIs, subsequently annotating the epithelial from the stromal compartment (45). Cell segmentation was performed using Qupath, extracting the mean of pixel intensities of each subchannel within each cellular segment, and x,y cell centroids. Spatially resolved single-cell data was exported as .csv. In, python, we limited the outlier signal to the [Quartile 3 + 1.5x(Inter Quartile Range)] and rescaled it to the [0,1] range using the MIN-MAX values for each marker. Stromal cell distance to nearest epithelial cell was assigned using an adaptation of our Mask-Constrained Nearest Neighbor Algorithm. Data analysis and plotting was performed in R, restricted to stromal cells located within 100µm from the epithelium.

### Cell lines and human primary fibroblasts

Primary fibroblast lines were established from patient #3870, #4156, and #3910 with pancreatic ductal adenocarcinoma enrolled in the hepatobiliary and pancreatic biobank at the CRCHUM under Dr. Simon Turcotte. Tumour specimens were transported on ice in transfer medium (**Supplementary Table 6**) and processed the same day under sterile conditions. Tissue was minced and cultured in fibroblast growth medium to allow outgrowth. Media was refreshed twice weekly until ∼70% confluency. Cells were passaged using 0.05% trypsin (3 min) and centrifuged at 300 × g for 10 min at 4 °C before expansion at a 1:2 split ratio. SU8686 (#CRL-1837) and PANC-1 (#CRL-1469) were obtained from ATCC (Cedarlane) and maintained according to manufacturer recommendations. To generate fluorescently labelled SU8686 and PANC-1 cell line, mCherry coding sequences were cloned into the dCAS9-VP64 backbone (Addgene, cat# 61422) in place of the dCAS9-VP64 insert to make lentivirus encoding the fluorophore under the EF1α promoter. Lentiviral particles were produced in HEK293T cells by calcium phosphate co-transfection of the transfer plasmid with the second-generation packaging plasmids psPAX2 (Addgene, cat# 12260) and pMD2.G (Addgene, cat# 12259) in 10 cm dishes. Viral supernatant was harvested at 72 hours post-transfection, filtered through a 0.45 µm membrane, and concentrated overnight by PEG/NaCl precipitation at 4°C. Target cells were transduced by reverse transduction, whereby 1 × 10⁵ cells resuspended in 30 µl of medium plus polybrene 0.4 µL/mL (Sigma, cat# TR-1003-G) were combined with 1 µl of concentrated lentiviral suspension directly in a 6-well plate, followed by addition of 2 ml of complete medium refreshed after 24 hours. Stably transduced cells were selected by fluorescence-activated cell sorting (FACS) of live mCherry-positive single cells on a BD FACSAria IIIu (4-laser configuration) at the IRIC Flow Cytometry Core Facility 72 hours post-transduction. For 3D assays, cells were adapted to conditioned growth medium for two weeks prior to the seeding date (**Supplementary Table 6**). All cultures were maintained at 37 °C, 5% CO₂.

### 3D tumoroid–fibroblast co-culture

Tumoroids were generated in 96-well AggreWell plates by seeding 100 tumour cells per tumoroid (32 tumoroids/well) in 200 µL medium and incubating overnight. On Day 2, fibroblasts (500 cells per tumoroid) were added after removal of 100 µL medium. On Day 3, tumoroids were collected, transferred to 96-well imaging plates, and embedded in 33% Matrigel (15 µL Matrigel + cell suspension). Gels were polymerized for 20–30 min at 37 °C, followed by addition of complete medium and baseline imaging. Treatment was initiated on Day 4 (2× dose as indicated, 200 µL). A first boost (1×) was applied on Day 7 by replacing 100 µL medium with fresh treatment, followed by a second identical boost on Day 9. Cultures were maintained until Day 18 for final imaging and endpoint analysis. Imaging was performed on a Cytation 5 imaging reader (BioTek) at 4× magnification, acquiring DAPI (nuclei), Texas Red (tumor cells), and phase contrast channels. Images were acquired as 21 optical sections at a Z-step of 10 µm.

Custom image analysis was performed. Z-stack images were processed in Fiji using a custom macro. For each Texas Red image stack, a maximum-intensity projection was computed across all slices using the Z Project command (projection type: Max Intensity), collapsing the stack into a single two-dimensional image and saving it as a TIFF. To visualize temporal dynamics, maximum intensity projections from each well were color-coded by timepoint using a custom shell script based on ImageMagick (single common intensity range (minimum and maximum) was computed and applied across all timepoints of a given well). Composite images per well were generated by overlaying all timepoints with the earliest in the foreground. Single spheroid segmentation, tracking over time, and morphometric quantification (tumor cell proliferation and organoid circularity) were performed in Qupath (45) and Python from Texas Red fluorescent images. From raw images, positive pixels were defined by pixel classification using thresholding. Single spheroid tracking was performed using a custom groovy code, where positive regions were merged and separated into connected components, and components below 3,000 px² were discarded as debris. At the baseline timepoint (T0), each spheroid was assigned an identity and its centroid and morphology recorded. At each subsequent timepoint, positive components were matched to T0 spheroids by containment of the T0 centroids (components containing a single T0 centroid were assigned directly, components containing multiple T0 centroids were partitioned by Voronoi tessellation, and components containing no T0 centroid were assigned to the nearest T0 spheroid within 500 px or discarded. For all tracked spheroids, we computed projected area, perimeter, and shape descriptors, including circularity, with per-spheroid changes relative to T0. Data analysis and plotting were performed in Python.

### 2D tumor–fibroblast co-culture

96-well plates were coated with 1% matrigel (Sigma, cat# CLS356231) for 30 min at 37°C, 5% CO2. After removing matrigel, co-cultures were seeded at a 4:1 CAF-to-tumor ratio. Briefly, 2000 CAFs (primary cell lines 3870, 3910, 4156) and 500 mCherry-expressing SU8686 cells were plated per well and cultured for 5 days. Cells were maintained in DMEM/F-12 with GlutaMAX (Gibco, Thermo Fisher Scientific, cat# 10565018) supplemented with 1× N-2 (Gibco, Thermo Fisher Scientific, cat# 17502048) and 1× B-27 supplement (Gibco, Thermo Fisher Scientific, cat# 17504044). Medium was refreshed every other day. Control conditions included SU8686 cells cultured alone (tumor growth control), CAFs cultured alone in the same medium as the co-culture (negative control for marker expression). After 5 days of culture, cells were fixed with 4% paraformaldehyde (PFA; Electron Microscopy Sciences/Cedarlane, cat# 15710) in PBS at 37°C for 30 minutes, followed by two washes with PBS. Permeabilization was performed with 0.1% Tween-20 (BioShop, cat# TWN510.100) in PBS supplemented with 2% FBS for 10 minutes at room temperature. Cells were then incubated overnight at 4°C with an Alexa Fluor 594 anti-Nestin antibody (clone 10C2; BioLegend, cat# 656804, dilution 1:2000) diluted in PBS + 2% FBS + 0.1% Tween-20. Following three washes with PBS + 2% FBS, cells were incubated with DAPI (0.1 µg/ml; Sigma-Aldrich, cat# D9542) for nuclear counterstaining. Imaging was performed on a Cytation 5 imaging reader (BioTek) at 4× magnification, acquiring DAPI (nuclei), Texas Red (NESTIN), and phase contrast channels. Custom image analysis was performed in FIJI (v1.54), Ilastik (v1.4.0), and CellProfiler (v4.2.6). Raw images underwent illumination correction by rolling-ball background subtraction (radius = 50 pixels) in FIJI. Two brightness-corrected DAPI images were generated per well: one optimized for nuclear segmentation, and an over-saturated version in which the diffuse cytoplasmic DAPI signal outlines the whole-cell body of both CAFs and tumor cells. DAPI nuclei were identified as primary objects in CellProfiler using intensity-based declumping. Whole-cell body objects were propagated from these nuclei using an Ilastik pixel-classification probability map trained on the over-saturated DAPI image. The mCherry (Texas Red) channel was separately used in Ilastik to generate a tumor-specific probability map, from which tumor nuclei and tumor body objects were identified in CellProfiler. Each whole-cell body object was classified as tumor or CAF based on overlap with this tumor object set (FilterObjects/RelateObjects), and individual binary CAF and tumor masks were regenerated from the classified objects. Masks were exported as TIFF images, and per-object shape and marker intensity measurements were exported to CSV via ExportToSpreadsheet. For each co-culture well, a reference image was reconstructed from the segmentation masks, assigning CAFs (Population A) and Su8686 tumor cells (Population B) distinct colors; CAF identities were recovered from a per-cell label image built by mapping each segmented mask pixel to its nearest CellProfiler object centroid. Marker intensities were normalized per well to a 0–1 range for visualization, with the 2nd and 98th percentiles set as the minimum and maximum to reduce outlier influence. Fibroblast distance to the nearest tumor cell was assigned using an adaptation of our Mask-Constrained Nearest Neighbor Algorithm.

## Ethics board approval

The project was completed under main project 2026-6334. The computation pathology component and image repository access is under 2025-12351. The primary line generation was performed under project 23.031.

## Data availability

WSIs, CyCIF images, and **Supplementary Table 1** are available by contacting the corresponding author and establishing an academic material transfer agreement. Processed CyCIF data, spatial transcriptomic data, **Supplementary Table 4**, and **Supplementary Table 5** are freely available through Open Science Framework (osf.io/yd8zh).

## Code availability

All code will be made publicly available on GitHub upon peer-reviewed publication.

## ACKNOWLEDGMENTS

We thank Wiliam Renock from the Yale Center for Genomic Analysis Core for Xenium *in situ* imaging processing. We thank Frank Revetta for preparing FFPE samples. We thank Mélina Narlis and Julie Hinsinger from the Institute for Research in Immunology and Cancer Research Histology Core. M.B. was supported by Fonds Pierre-Saul. V.Q.T. is supported by the Fonds de recherche Québec Santé J1 Clinical-Scholar program, the Canadian Cancer Society Breakthrough Grant, Amazon Science, Institute for Research in Immunology start-up funds, the Canadian Foundation for Innovation John Evans Leaders Fund, the Digital Research Alliance Canada Research Allocation Competition, the Chan-Soon-Shiong Family Foundation, and the Université de Montréal salary support for clinician-scientists. D.J.H.F.K. is supported by the Fonds de recherche Québec Santé J1 283502. S.F.R. acknowledges funding from the Terry Fox Research Institute, Fonds de Recherche en Santé du Québec and V Foundation for Cancer Research.

## AUTHOR CONTRIBUTIONS

**M. Batardière:** Conceptualization, data curation, formal analysis, investigation, methodology, software, visualization, validation, writing – original draft, writing – review & editing**. M.I. Ibanez-Rios:** Conceptualization, data curation, formal analysis, investigation, methodology, software, visualization, validation, writing – review & editing. **A. Khellaf:** Conceptualization, data curation, formal analysis, investigation, methodology, software, visualization, validation, writing – review & editing. **E. Dianati:** Conceptualization, formal analysis, investigation, methodology, project administration, resources, supervision, validation, writing – original draft, writing – review & editing**. S.S. Diwan:** data curation, investigation, formal analysis, methodology, software, writing – review & editing. **M. Pelloux:** Conceptualization, data curation, formal analysis, investigation, methodology, software, writing – review & editing. **A. Archambault-Marsan:** data curation, formal analysis, investigation, methodology, writing – review & editing. **C. Beaussier**: data curation, investigation, methodology, writing – review & editing. **J. Diwan:** data curation, investigation, methodology, software, writing – review & editing. **S. Sapon-Cousineau:** data curation, investigation, writing – review & editing. **J. Baig:** data curation, investigation, methodology, software, writing – review & editing. **A. Shoukari:** data curation, formal analysis, investigation, methodology, writing – review & editing. **Z. Ghanmeh:** data curation, methodology, software, writing – review & editing. **Z. Yang:** data curation, methodology, software, writing – review & editing. **M. Farias Gonzalez:** data curation, investigation, methodology, software, writing – review & editing. **J.L. Li:** data curation, methodology, software, writing – review & editing. **A. Kessab:** data curation, methodology, software, writing – review & editing. **L. Lando:** Conceptualization, writing – review & editing. **P. Lefrancois:** Conceptualization, writing – review & editing. **S. Turcotte:** Conceptualization, resources, writing – review & editing. **Y. A. Lee:** Conceptualization, writing – review & editing. **S.F. Roy:** Conceptualization, resources, writing – review & editing. **M.S. Hosseini:** Conceptualization, writing – review & editing. **B. Tessier-Cloutier:** Conceptualization, resources, writing – review & editing. **M.C.B. Tan:** Conceptualization, writing – review & editing. **K.E. DelGiorno:** Conceptualization, writing – review & editing. **D.J.H.F. Knapp:** Conceptualization, data curation, formal analysis, investigation, methodology, project administration, resources, software, supervision, validation, visualization, writing – review & editing. **V.Q. Trinh:** Conceptualization, data curation, formal analysis, funding acquisition, investigation, methodology, project administration, resources, supervision, validation, writing – original draft, writing – review & editing.

## SUPPLEMENTAL TABLE LEGENDS

**Table S1. Clinical data.** Clinical data for computational pathology pipeline, CyCIF, and Xenium *In Situ* cohorts.

**Table S2. CyCIF Supplemental data.** Tables of antibody panel information and sample region of interest distribution.

**Table S3.** Xenium custom gene panel list.

**Table S4. Top differentially expressed genes per population.** Tables including top differentially expressed genes and LISI scores for Crude clusters, and for each Refined cluster.

**Table S5. Ligand-receptor analysis outputs.**

**Table S6. Supplemental material and methods for in vitro assay.**

## SUPPLEMENTAL FIGURE LEGENDS

**Figure S1. Additional information for computational pathology analysis. A**. Table of histomorphology semantic characterization of clusters from computational pathology pipeline. **B.** Stacked bar plot showing the proportion of each cluster per grade (stromal vs epithelial). **C.** Table summarizing ROI count per cluster ID and per grade. **D**. Bar plots of proportion of each cluster per grade (one plot per cluster).

**Figure S2. Epithelial-stromal distances analysis of immune and endothelial cells. A**. Connected dot plot of proportions of immune and endothelial cells sub-populations as a function of cell distance to nearest epithelial cell.

**Figure S3. Epithelial-stromal distances analysis of stromal content relative to adjacent epithelial cell identity. A**. Schematic overview of our radius search-based distance analyses of fibroblasts and immune cells, showing cells detected within 50µm of epithelial cells in green, and those outside of distance range in red. **B**. Proportion of fibroblast clusters relative to adjacent epithelial subtypes. **C**. Proportion of immune clusters relative to adjacent epithelial subtypes.

**Figure S4. Representative Xenium in situ quality control metrics cLISI and iLISI.** UMAP projections colored by sample (**A**), Crude cluster (**B**), cLISI (**C**), and iLISI (**D**).

**Figure S5. Improved fibroblast detection with anucleated segmentation pipeline. A.** Representation of additionally segmented cells with anucleated pipeline, showing unassigned RNA, and RNA assigned to a cell segment (anucluated or nucleated). **B.** Proportion of anucleated cells per Crude Cluster.

**Figure S6. Top cell populations of SpaTopic analysis.** Matrix plot showing the proportion of each sub-population per topic.

**Figure S7. Giotto co-localization analysis.** Heatmap of cell population co-localisation analysis of LG, HG, and INV respectively.

**Figure S8. Ligand-receptor predictions of SpaTopic 6 cell populations.** Ligand-receptor network between cell types for each LG, HG, and INV grades, cells of SpaTopic 6 only.

## REFERENCES

1. American Cancer Society. Cancer Facts & Figures 2025. [Available from: https://www.cancer.org/content/dam/cancer-org/research/cancer-facts-and-statistics/annual-cancer-facts-and-figures/2025/2025-cancer-facts-and-figures-acs.pdf.

2. Siegel RL, Giaquinto AN, Jemal A. Cancer statistics, 2024. CA Cancer J Clin. 2024;74(1):12–49.

3. Li S, Xie K. Ductal metaplasia in pancreas. Biochim Biophys Acta Rev Cancer. 2022;1877(2):188698.

4. Diana Agostini-Vulaj DO. Pancreas Cystic and intraductal lesions Intraductal papillary mucinous neoplasm (IPMN) 2021 [Available from: https://www.pathologyoutlines.com/topic/pancreasipmn.html.

5. Makino Y, Oyama K, Sagara A, Thege FI, Maitra A. Molecular pathology of intraductal papillary mucinous neoplasms of the pancreas: current understanding and perspectives on malignant progression. J Gastroenterol. 2026;61(7):909–24.

6. Pea A, He X, Upstill-Goddard R, Luchini C, Schubert Santana LP, Dreyer S, et al. Clonal evolutionary analysis reveals patterns of malignant transformation of Intraductal Papillary Mucinous Neoplasms of the pancreas. Nat Commun. 2026;17(1).

7. Li J, Branch G, Macchia J, Elhossiny AM, Arya N, Liang J, et al. Spatial Analysis of Intraductal Papillary Mucinous Neoplasms Defines a Paradoxical Keratin 17-Positive, Low-Grade Epithelial Population Harboring Malignant Features. Cell Mol Gastroenterol Hepatol. 2026;20(6):101749.

8. Cui M, Mo S, Bai J, Javed AA, Habib JR, Yang S, et al. Spatial transcriptomics defines the molecular progression, invasion and immune landscape of IPMN and IPMN-derived pancreatic cancer. Gut. 2026;75(4):801–14.

9. Huang X, Feng Y, Ma D, Ding H, Dong G, Chen Y, et al. The molecular, immune features, and risk score construction of intraductal papillary mucinous neoplasm patients. Front Mol Biosci. 2022;9:887887.

10. Ma Z, Lytle NK, Chen B, Jyotsana N, Novak SW, Cho CJ, et al. Single-Cell Transcriptomics Reveals a Conserved Metaplasia Program in Pancreatic Injury. Gastroenterology. 2022;162(2):604–20 e20.

11. José Reyes IDP, Andrea C. Chaikovsky, Nikhita Pasnuri, Ahmed M. Elhossiny, Jin Park, Philipp Weiler, Tobias Krause, Andrew Moorman, Catherine Snopkowski, Meril Takizawa, Cassandra Burdziak, Nalin Ratnayeke, Ignas Masilionis, Yu-Jui Ho, Ronan Chaligné, Paul B. Romesser, Aveline Filliol, Tal Nawy, John P. Morris, Zhen Zhao, Marina Pasca Di Magliano, Direna Alonso-Curbelo, Dana Pe’er, Scott W. Lowe. Oncogenic and tumor-suppressive forces converge on a progenitor niche at the benign-to-malignant transition. Cell. 2026;189(10).

12. Quoc-Huy Trinh V, Ankenbauer KE, Torbit SM, Taranto CP, Liu J, Batardiere M, et al. Mutant GNAS drives a pyloric metaplasia with tumor suppressive glycans in intraductal papillary mucinous neoplasia. Cell Rep. 2025;44(12):116684.

13. Wang Q, Shao X, Zhang Y, Zhu M, Wang FXC, Mu J, et al. Role of tumor microenvironment in cancer progression and therapeutic strategy. Cancer Med. 2023;12(10):11149–65.

14. Li J, Chen D, Shen M. Tumor Microenvironment Shapes Colorectal Cancer Progression, Metastasis, and Treatment Responses. Front Med (Lausanne). 2022;9:869010.

15. Akinsipe T, Mohamedelhassan R, Akinpelu A, Pondugula SR, Mistriotis P, Avila LA, Suryawanshi A. Cellular interactions in tumor microenvironment during breast cancer progression: new frontiers and implications for novel therapeutics. Front Immunol. 2024;15:1302587.

16. Jiao F, Wang Z, Yuan J, Shi F, Zhang S. The tumor microenvironment shapes gastric cancer progression by coordinating immune suppression and metabolic reprogramming. Front Immunol. 2026;17:1787060.

17. Kwon JY, Vera RE, Fernandez-Zapico ME. The multi-faceted roles of cancer-associated fibroblasts in pancreatic cancer. Cell Signal. 2025;127:111584.

18. Hartupee C NB, Chabu CY, Tesfay MZ, Coleman-Barnett J, West JT and Moaven O. Pancreatic cancer tumor microenvironment is a major therapeutic barrier and target. Front Immunol. 2024;15.

19. Sunami Y, Boker V, Kleeff J. Targeting and Reprograming Cancer-Associated Fibroblasts and the Tumor Microenvironment in Pancreatic Cancer. Cancers (Basel). 2021;13(4).

20. Brichkina A, Polo P, Sharma SD, Visestamkul N, Lauth M. A Quick Guide to CAF Subtypes in Pancreatic Cancer. Cancers. 2023;15(9):2614.

21. Zhang T, Ren Y, Yang P, Wang J, Zhou H. Cancer-associated fibroblasts in pancreatic ductal adenocarcinoma. Cell Death Dis. 2022;13(10):897.

22. Huang H, Brekken RA. Recent advances in understanding cancer-associated fibroblasts in pancreatic cancer. Am J Physiol Cell Physiol. 2020;319(2):C233–C43.

23. Giulia Biffi TEO, Benjamin Spielman, Yuan Hao, Ela Elyada, Youngkyu Park, Jonathan Preall, David A Tuveson. IL-1-induced JAK/STAT signaling is antagonized by TGF-β to shape CAF heterogeneity in pancreatic ductal adenocarcinoma. Cancer Discov. 2020.

24. Ozdemir BC, Pentcheva-Hoang T, Carstens JL, Zheng X, Wu CC, Simpson TR, et al. Depletion of Carcinoma-Associated Fibroblasts and Fibrosis Induces Immunosuppression and Accelerates Pancreas Cancer with Reduced Survival. Cancer Cell. 2015;28(6):831–3.

25. Ohlund D, Handly-Santana A, Biffi G, Elyada E, Almeida AS, Ponz-Sarvise M, et al. Distinct populations of inflammatory fibroblasts and myofibroblasts in pancreatic cancer. J Exp Med. 2017;214(3):579–96.

26. Bernard V, Semaan A, Huang J, San Lucas FA, Mulu FC, Stephens BM, et al. Single-Cell Transcriptomics of Pancreatic Cancer Precursors Demonstrates Epithelial and Microenvironmental Heterogeneity as an Early Event in Neoplastic Progression. Clinical Cancer Research. 2019;25(7):2194–205.

27. Li J, Wei T, Ma K, Zhang J, Lu J, Zhao J, et al. Single-cell RNA sequencing highlights epithelial and microenvironmental heterogeneity in malignant progression of pancreatic ductal adenocarcinoma. Cancer Lett. 2024;584:216607.

28. Iyer MK, Fletcher AA, Okoye JO, Shi C, Chen F, Kanu EN, et al. Spatial Transcriptomics of Intraductal Papillary Mucinous Neoplasms Reveals Divergent Indolent and Malignant States. Clin Cancer Res. 2025;31(9):1796–808.

29. Enzler T, Shi J, McGue J, Griffith BD, Sun L, Sahai V, et al. A Comparison of Spatial and Phenotypic Immune Profiles of Pancreatic Ductal Adenocarcinoma and Its Precursor Lesions. Int J Mol Sci. 2024;25(5).

30. Shih AH, Holland EC. Notch signaling enhances nestin expression in gliomas. Neoplasia. 2006;8(12):1072–82.

31. Hernandez S, Parra ER, Uraoka N, Tang X, Shen Y, Qiao W, et al. Diminished Immune Surveillance during Histologic Progression of Intraductal Papillary Mucinous Neoplasms Offers a Therapeutic Opportunity for Cancer Interception. Clin Cancer Res. 2022;28(9):1938–47.

32. Jamouss KT, Damanakis AI, Cornwell AC, Jongepier M, Trujillo MA, Pflüger MJ, et al. Tumor immune microenvironment alterations associated with progression in human intraductal papillary mucinous neoplasms. Journal of Pathology. 2025;266(1):40–50.

33. Bell ATF, Mitchell JT, Kiemen AL, Lyman M, Fujikura K, Lee JW, et al. PanIN and CAF transitions in pancreatic carcinogenesis revealed with spatial data integration. Cell Syst. 2024;15(8):753–69 e5.

34. Elhossiny AM, Kadiyala P, Okoye JO, Hiraki HL, Procario MC, Giridharan T, et al. Asynchronous evolution of epithelium and stroma differentiates precursor lesions from pancreatic cancer. Cancer Discov. 2026.

35. Peng S, Chen Q, Chen Z, Yao M, Cai Y, He D, et al. Evolution of the Spatial transcriptomic landscape during the progression of high-grade pancreatic intraductal papillary mucinous neoplasms to invasive cancer. Pancreatology. 2026;26(2):246–58.

36. Nielsen MFB, Mortensen MB, Detlefsen S. Typing of pancreatic cancer-associated fibroblasts identifies different subpopulations. World J Gastroenterol. 2018;24(41):4663–78.

37. Tong Z, Yin Z. Distribution, contribution and regulation of nestin(+) cells. J Adv Res. 2024;61:47–63.

38. Zhou B, Lin W, Long Y, Yang Y, Zhang H, Wu K, Chu Q. Notch signaling pathway: architecture, disease, and therapeutics. Signal Transduct Target Ther. 2022;7(1):95.

39. Ghosh A, Mitra AK. Metastasis and cancer associated fibroblasts: taking it up a NOTCH. Front Cell Dev Biol. 2023;11:1277076.

40. Shinyoung Kim KY, Kiyoon Kim, Hee Ja Kim, Da Young Kim, Jeesoo Chae, Young-Ho Ahn, Jihee Lee Kang. The interplay of cancer-associated fibroblasts and apoptotic cancer cells suppresses lung cancer cell growth through WISP-1-integrin ανβ3-STAT1 signaling pathway. Cell Communication and Signaling. 2025;23.

41. Du Y, Shao H, Moller M, Prokupets R, Tse YT, Liu ZJ. Intracellular Notch1 Signaling in Cancer-Associated Fibroblasts Dictates the Plasticity and Stemness of Melanoma Stem/Initiating Cells. Stem Cells. 2019;37(7):865–75.

42. Hongwei Shao MM, Long Cai, Rochelle Prokupets, Cuixia Yang, Connor Costa, Kerstin Yu, Nga Le, Zhao-Jun Liu. Converting melanoma-associated fibroblasts into a tumor-suppressive phenotype by increasing intracellular Notch1 pathway activity. PLoS ONE. 2021.

43. Wang J, Lai X, Yao S, Chen H, Cai J, Luo Y, et al. Nestin promotes pulmonary fibrosis via facilitating recycling of TGF-beta receptor I. Eur Respir J. 2022;59(5).

44. Chandler C, Liu T, Buckanovich R, Coffman LG. The double edge sword of fibrosis in cancer. Transl Res. 2019;209:55–67.

45. Bankhead P, Loughrey MB, Fernandez JA, Dombrowski Y, McArt DG, Dunne PD, et al. QuPath: Open source software for digital pathology image analysis. Sci Rep. 2017;7(1):16878.

46. Martin Weigert US. Nuclei instance segmentation and classification in histopathology images with StarDist. arXiv. 2022(Computer Vision and Pattern Recognition).

47. Chen RJ, Ding T, Lu MY, Williamson DFK, Jaume G, Chen B, et al. A General-Purpose Self-Supervised Model for Computational Pathology. ArXiv. 2023.

48. Maxime Oquab TD, Théo Moutakanni, Huy Vo, Marc Szafraniec, Vasil Khalidov, Pierre Fernandez, Daniel Haziza, Francisco Massa, Alaaeldin El-Nouby, Mahmoud Assran, Nicolas Ballas, Wojciech Galuba, Russell Howes, Po-Yao Huang, Shang-Wen Li, Ishan Misra, Michael Rabbat, Vasu Sharma, Gabriel Synnaeve, Hu Xu, Hervé Jegou, Julien Mairal, Patrick Labatut, Armand Joulin, Piotr Bojanowski. DINOv2: Learning Robust Visual Features without Supervision. arXiv. 2023(Computer Vision and Pattern Recognition).

49. Adam Paszke SG, Francisco Massa, Adam Lerer, James Bradbury, Gregory Chanan, Trevor Killeen, Zeming Lin, Natalia Gimelshein, Luca Antiga, Alban Desmaison, Andreas Köpf, Edward Yang, Zach DeVito, Martin Raison, Alykhan Tejani, Sasank Chilamkurthy, Benoit Steiner, Lu Fang, Junjie Bai, Soumith Chintala. PyTorch: An Imperative Style, High-Performance Deep Learning Library. arXiv. 2019(Machine Learning).

50. Bahman Bahmani BM, Andrea Vattani, Ravi Kumar, Sergei Vassilvitskii. Scalable K-Means++. arXiv. 2012(Databases).

