## Supplementary Fig. S1 for "Spatial multi-omics resolve epithelium-fibroblast gradients and highlight NESTIN-NOTCH1-expressing subepithelial fibroblasts during human pancreatic tumorigenesis"

A

| Cluster | Semantic characterization | Code (Epithelial/ Stromal/ Other) | If Epithelial: Consistently atypical? |
|---|---|---|---|
| 0 | Stromal: stroma with lymphoplasmacytic infiltration | Stromal | |
| 1 | Stromal: stromal fibroblasts with collagen and no to minimal inflammatory cells **out of focus** | Stromal | |
| 2 | Epithelial: benign pancreatic ductal epithelium (small ducts) with surrounding stroma | Epithelial | no |
| 3 | Stromal: stromal fibroblasts with well-organized collagen fibers and no to minimal immune cells (lymphocytes/mast cells) | Stromal | |
| 4 | Stromal: stromal fibroblasts with loose collagen and no to minimal immune cells | Stromal | |
| 5 | Epithelial: normal to low grade atypia epithelial cells with underlying basement membrane | Epithelial | no |
| 6 | Stromal: vascularized stroma with loose collagen | Stromal | |
| 7 | Stromal: desmoplastic stroma with occasional plasmocytic infiltration | Stromal | |
| 8 | Epithelial: epithelial mucinous cells with low-grade atypia | Epithelial | yes |
| 9 | Stromal: inflammatory stroma with lymphoplasmacytic predominance | Stromal | |
| 10 | Epithelial: normal acinar epithelial cells with or without surrounding stroma | Epithelial | no |
| 11 | Stromal: normocellular stroma with fibroblasts, small capillaries and occasional erythrocyte extravasation and disorderly collagen and no to minimal inflammatory cells | Stromal | |
| 12 | Epithelial: epithelial cells with high-grade atypia | Epithelial | yes |
| 13 | Epithelial: low-grade mucinous epithelium with surrounding underlying stroma | Epithelial | yes |
| 14 | Epithelial: surface/luminal epithelial cells with high-grade atypia | Epithelial | yes |
| 15 | Epithelial: epithelial mucin-rich cells with low-grade atypia | Epithelial | yes |
| 16 | Epithelial: epithelial cells with low-grade atypia | Epithelial | yes |
| 17 | Stromal: normocellular to hypercellular stroma with activated fibroblasts and occasional capillaries | Stromal | |
| 18 | Epithelial: benign normal epithelial cells | Epithelial | no |
| 19 | Stromal: stromal capillaries with surrounding collagen fibers and occasional stromal erythrocyte extravasation | Stromal | |
| 20 | Stromal: normocellular to hypercellular vascularized stroma with well-organized collagen fibers | Stromal | |
| 21 | Stromal: stroma with capillaries and turgescent endothelial cells and occasional fibroblast | Stromal | |
| 22 | Epithelial: epithelial cells with low-grade to high-grade atypia | Epithelial | yes |
| 23 | Epithelial: epithelial cells with low-grade to high-grade atypia | Epithelial | yes |
| 24 | Stromal: normocellular stroma with fibroblasts, collagen and no to minimal inflammatory cells | Stromal | |
| 25 | Epithelial: epithelial mucinous cells with low-grade atypia | Epithelial | yes |
| 26 | Epithelial: epithelial mucinous cells with low-grade atypia | Epithelial | yes |
| 27 | Stromal: normocellular to hypercellular vascularized stroma with well-organized collagen fibers | Stromal | |
| 28 | Stromal: desmoplastic stroma with inflammatory predominance | Stromal | |
| 29 | Stromal: hypercellular desmoplastic stroma with activated fibroblasts, immune cells and disorderly collagen fibers | Stromal | |
| 30 | Epithelial: benign normal epithelial cells | Epithelial | no |
| 31 | Stromal: desmoplastic stroma with occasional activated fibroblasts | Stromal | |
| 32 | Stromal: subepithelial stroma with fibroblasts, collagen, capillaries and occasional immune cells | Stromal | |
| 33 | Stromal: stromal fibroblasts with loose collagen and occasional small capillaries and immune cells | Stromal | |
| 34 | Epithelial: epithelial cells with high-grade atypia | Epithelial | yes |

B

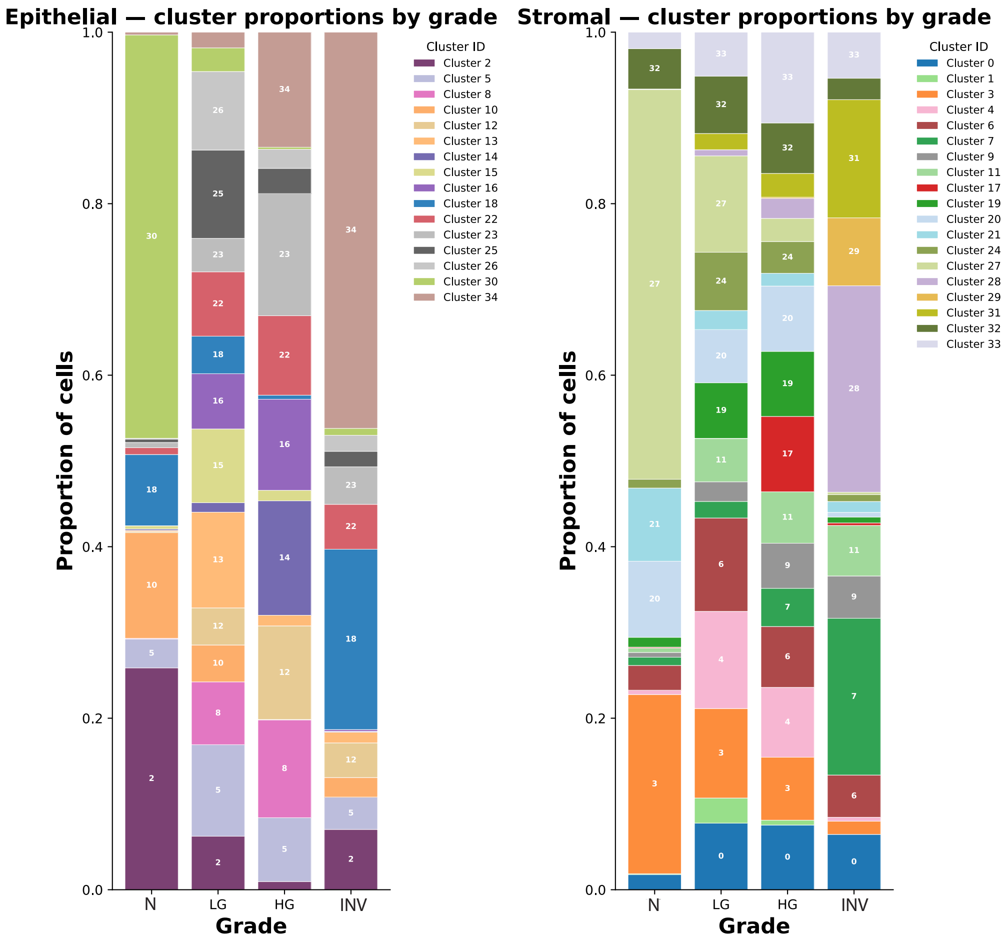


C

| cluster_id | N | LG | HG | INV | Total |
|---|---|---|---|---|---|
| 0 | 309 | 17488 | 12839 | 5757 | 36393 |
| 1 | 18 | 6576 | 967 | 0 | 7561 |
| 2 | 8925 | 8306 | 944 | 646 | 18821 |
| 3 | 3653 | 23483 | 12498 | 1373 | 41007 |
| 4 | 89 | 25503 | 13871 | 406 | 39869 |
| 5 | 1162 | 14221 | 7360 | 346 | 23089 |
| 6 | 504 | 24501 | 12080 | 4390 | 41475 |
| 7 | 168 | 4350 | 7583 | 16304 | 28405 |
| 8 | 27 | 9755 | 11277 | 1 | 21060 |
| 9 | 97 | 5145 | 8949 | 4398 | 18589 |
| 10 | 4253 | 5727 | 44 | 209 | 10233 |
| 11 | 90 | 11310 | 10216 | 5255 | 26871 |
| 12 | 43 | 5761 | 10790 | 371 | 16965 |
| 13 | 46 | 14864 | 1227 | 115 | 16252 |
| 14 | 57 | 1474 | 13197 | 8 | 14736 |
| 15 | 126 | 11453 | 1210 | 1 | 12790 |
| 16 | 4 | 8582 | 10497 | 19 | 19102 |
| 17 | 18 | 84 | 14963 | 264 | 15329 |
| 18 | 2865 | 5829 | 468 | 1932 | 11094 |
| 19 | 200 | 14593 | 12913 | 614 | 28320 |
| 20 | 1552 | 13969 | 12966 | 479 | 28966 |
| 21 | 1492 | 5006 | 2538 | 1135 | 10171 |
| 22 | 276 | 9991 | 9162 | 481 | 19910 |
| 23 | 208 | 5198 | 14079 | 402 | 19887 |
| 24 | 179 | 15280 | 6306 | 727 | 22492 |
| 25 | 132 | 13715 | 2908 | 166 | 16921 |
| 26 | 38 | 12196 | 2201 | 172 | 14607 |
| 27 | 7947 | 25297 | 4590 | 239 | 38073 |
| 28 | 7 | 1579 | 3983 | 21468 | 27037 |
| 29 | 0 | 49 | 236 | 7068 | 7353 |
| 30 | 16228 | 3676 | 239 | 74 | 20217 |
| 31 | 3 | 4239 | 4713 | 12291 | 21246 |
| 32 | 823 | 15041 | 10022 | 2222 | 28108 |
| 33 | 336 | 11554 | 18021 | 4794 | 34705 |
| 34 | 110 | 2437 | 13264 | 4245 | 20056 |
| Total | 51985 | 358232 | 269121 | 98372 | 777710 |

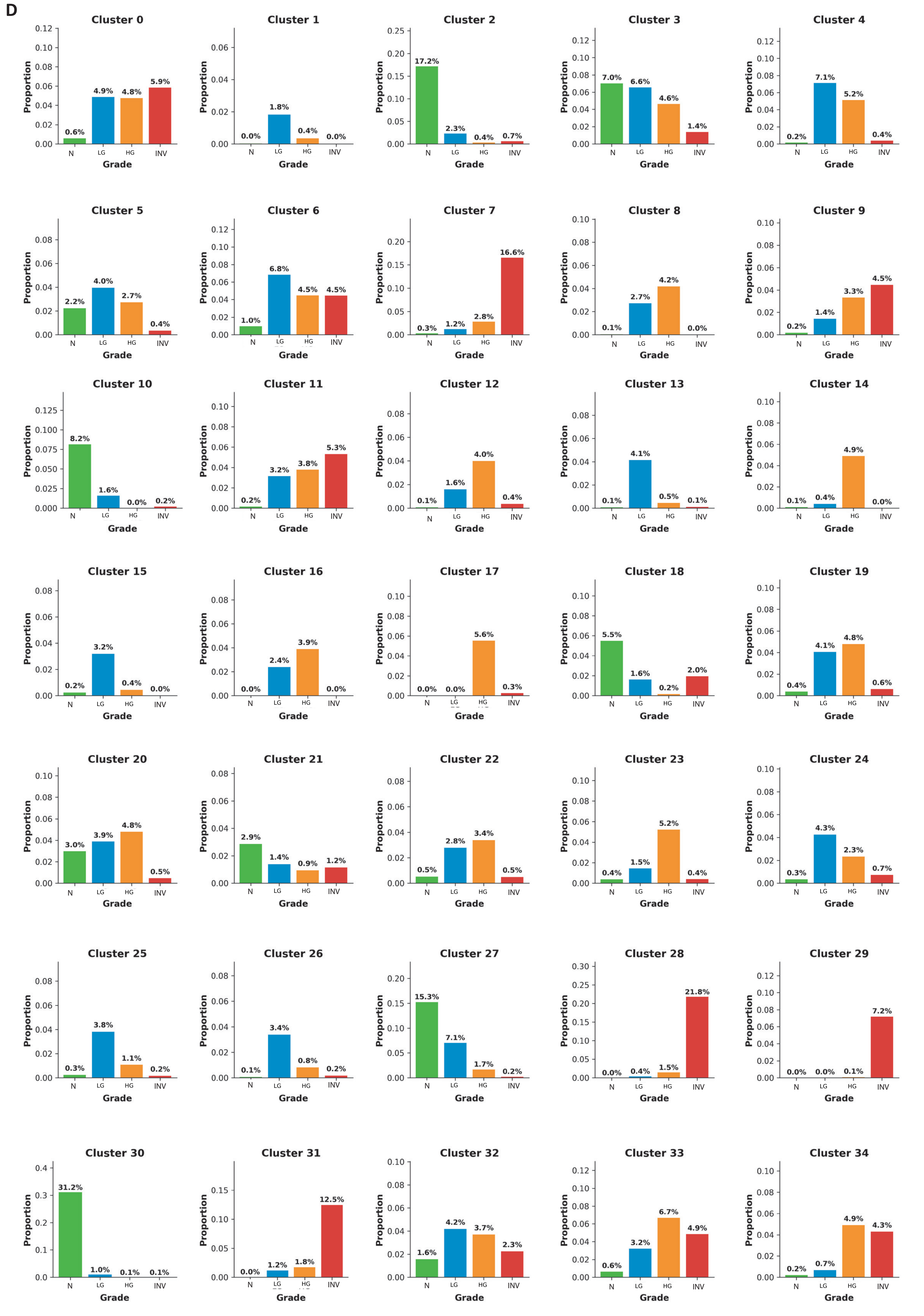
D
Cluster 0
0.6% 4.9% 4.8% 5.9%
Cluster 1
0.0% 1.8% 0.4% 0.0%
Cluster 2
17.2% 2.3% 0.4% 0.7%
Cluster 3
7.0% 6.6% 4.6% 1.4%
Cluster 4
0.2% 7.1% 5.2% 0.4%
Cluster 5
2.2% 4.0% 2.7% 0.4%
Cluster 6
1.0% 6.8% 4.5% 4.5%
Cluster 7
0.3% 1.2% 2.8% 16.6%
Cluster 8
0.1% 2.7% 4.2% 0.0%
Cluster 9
0.2% 1.4% 3.3% 4.5%
Cluster 10
8.2% 1.6% 0.0% 0.2%
Cluster 11
0.2% 3.2% 3.8% 5.3%
Cluster 12
0.1% 1.6% 4.0% 0.4%
Cluster 13
0.1% 4.1% 0.5% 0.1%
Cluster 14
0.1% 0.4% 4.9% 0.0%
Cluster 15
0.2% 3.2% 0.4% 0.0%
Cluster 16
0.0% 2.4% 3.9% 0.0%
Cluster 17
0.0% 0.0% 5.6% 0.3%
Cluster 18
5.5% 1.6% 0.2% 2.0%
Cluster 19
0.4% 4.1% 4.8% 0.6%
Cluster 20
3.0% 3.9% 4.8% 0.5%
Cluster 21
2.9% 1.4% 0.9% 1.2%
Cluster 22
0.5% 2.8% 3.4% 0.5%
Cluster 23
0.4% 1.5% 5.2% 0.4%
Cluster 24
0.3% 4.3% 2.3% 0.7%
Cluster 25
0.3% 3.8% 1.1% 0.2%
Cluster 26
0.1% 3.4% 0.8% 0.2%
Cluster 27
15.3% 7.1% 1.7% 0.2%
Cluster 28
0.0% 0.4% 1.5% 21.8%
Cluster 29
0.0% 0.0% 0.1% 7.2%
Cluster 30
31.2% 1.0% 0.1% 0.1%
Cluster 31
0.0% 1.2% 1.8% 12.5%
Cluster 32
1.6% 4.2% 3.7% 2.3%
Cluster 33
0.6% 3.2% 6.7% 4.9%
Cluster 34
0.2% 0.7% 4.9% 4.3%
Proportion
N LG HG INV
Grade
