## Supplementary figures and images for "Spatial multi-omics resolve epithelium-fibroblast gradients and highlight NESTIN-NOTCH1-expressing subepithelial fibroblasts during human pancreatic tumorigenesis"

### Supplementary Fig. S2

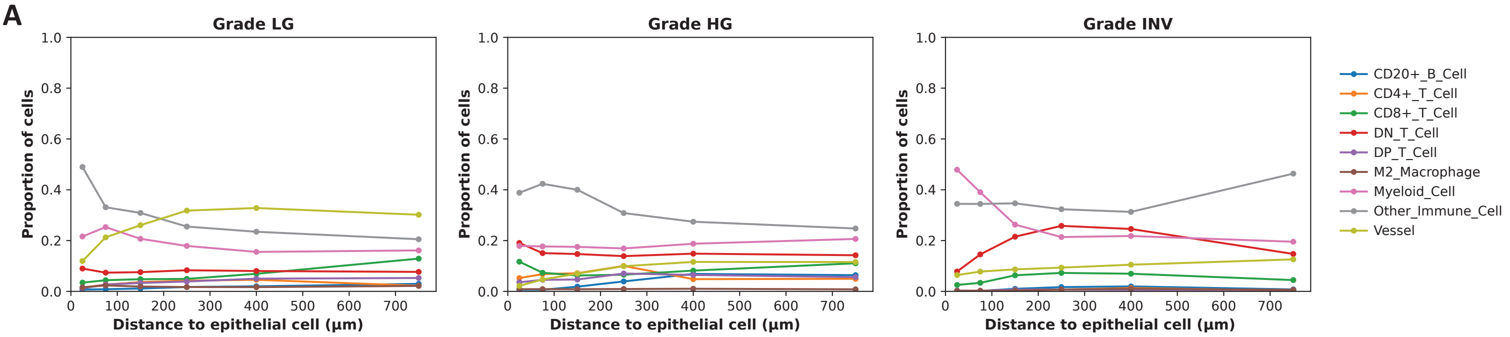

### Supplementary Fig. S3

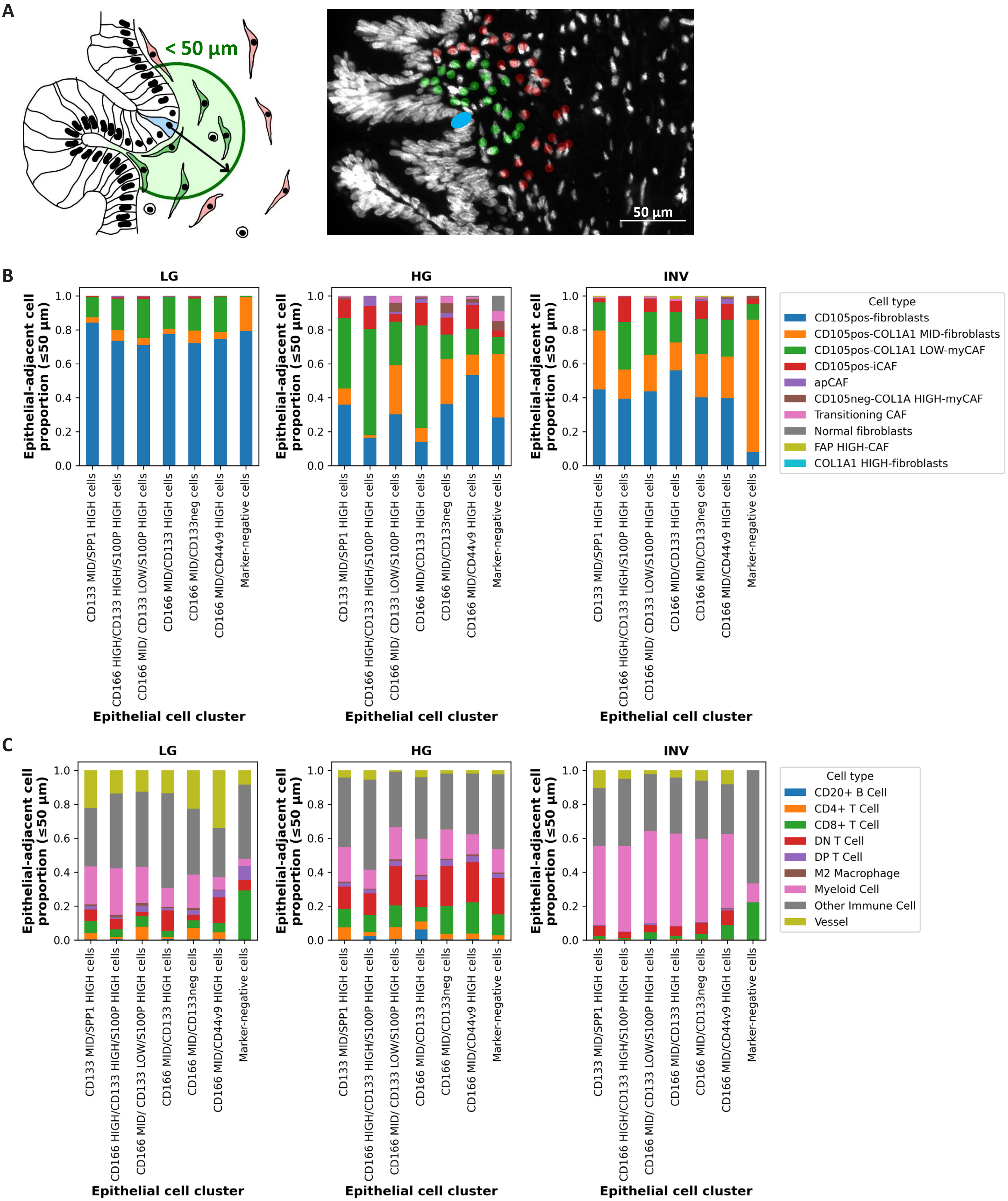

### Supplementary Fig. S4

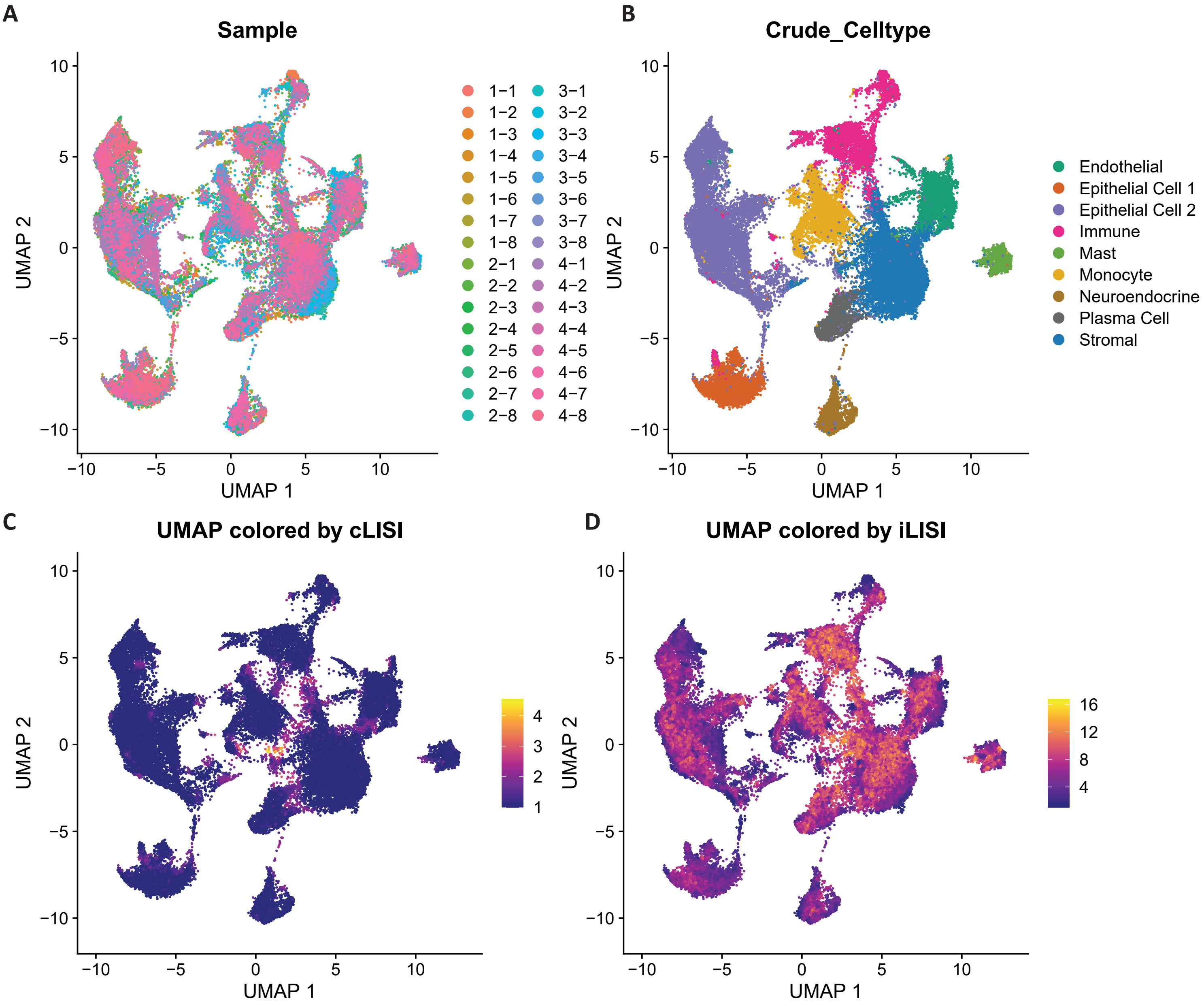

### Supplementary Fig. S5

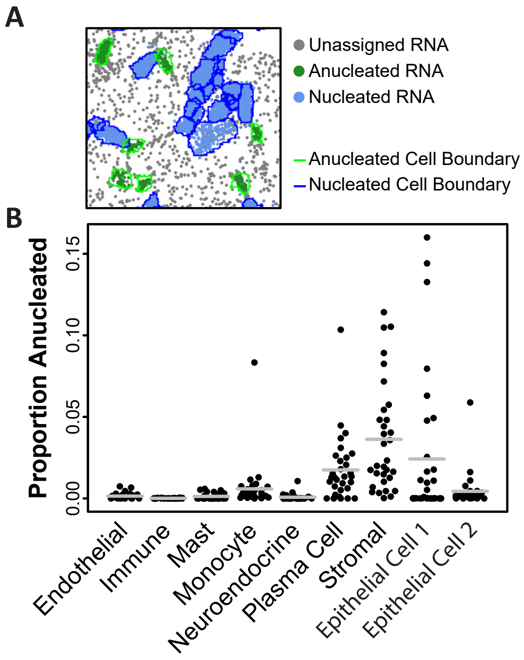

### Supplementary Fig. S6

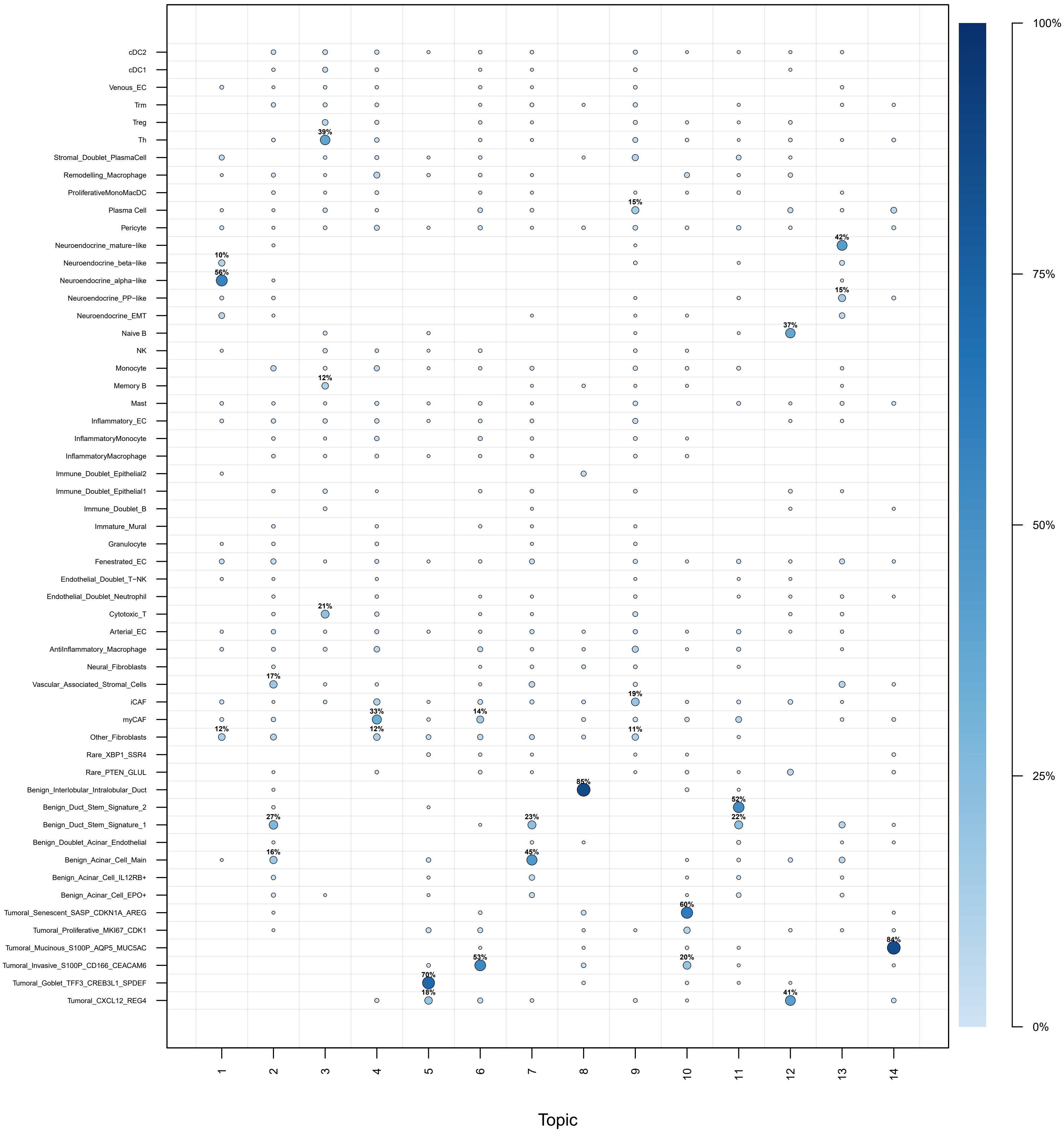

### Supplementary Fig. S7

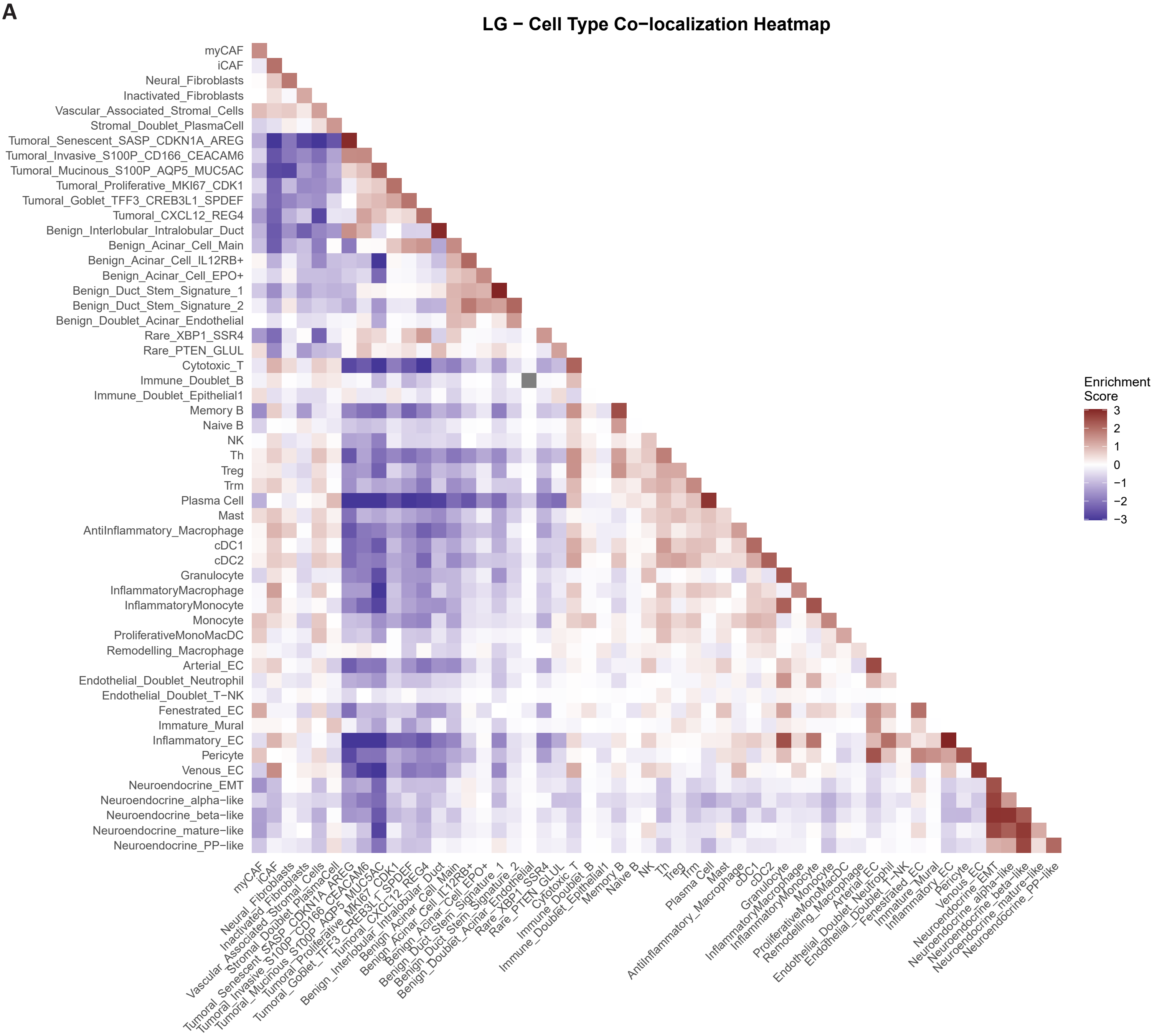

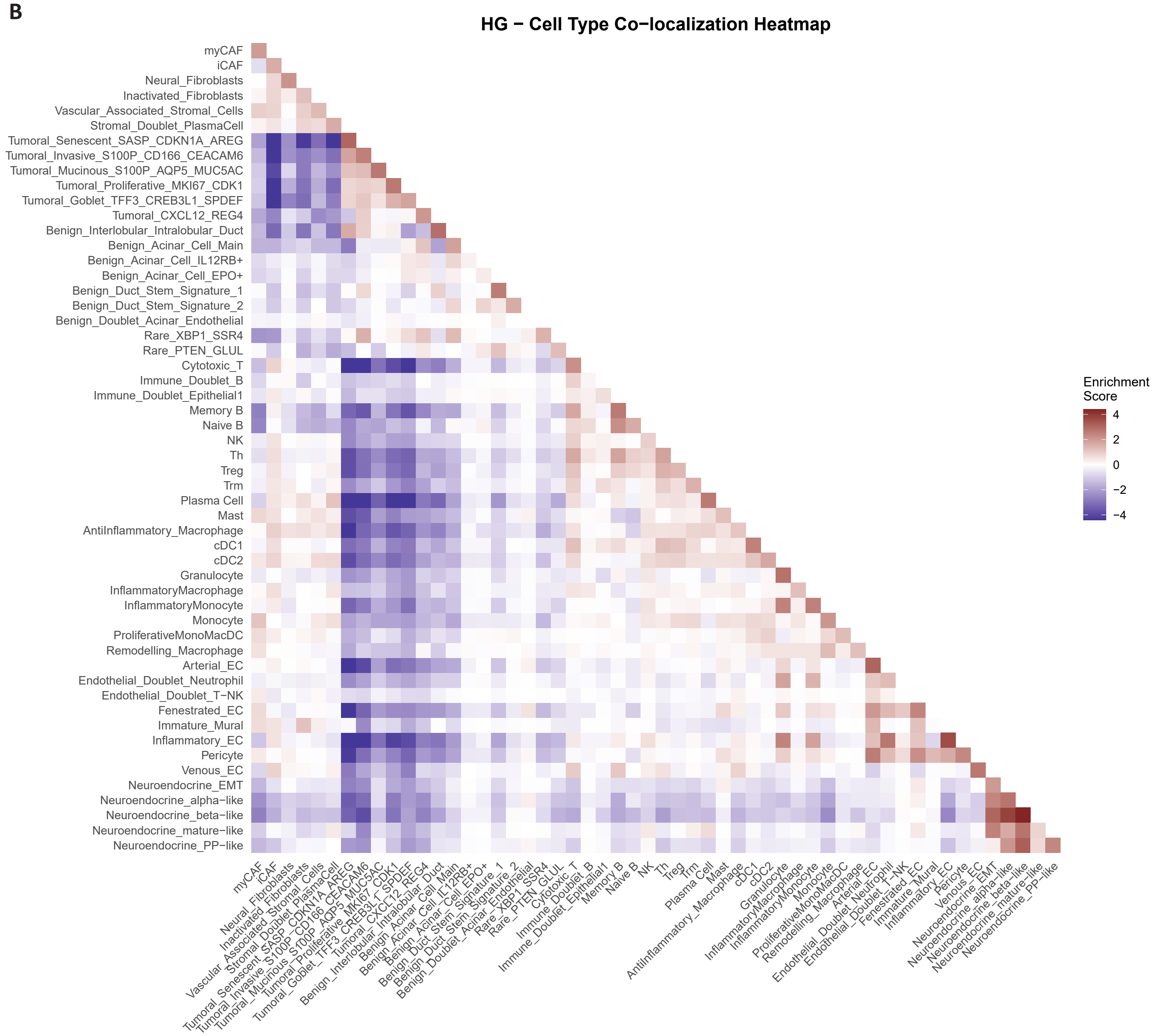

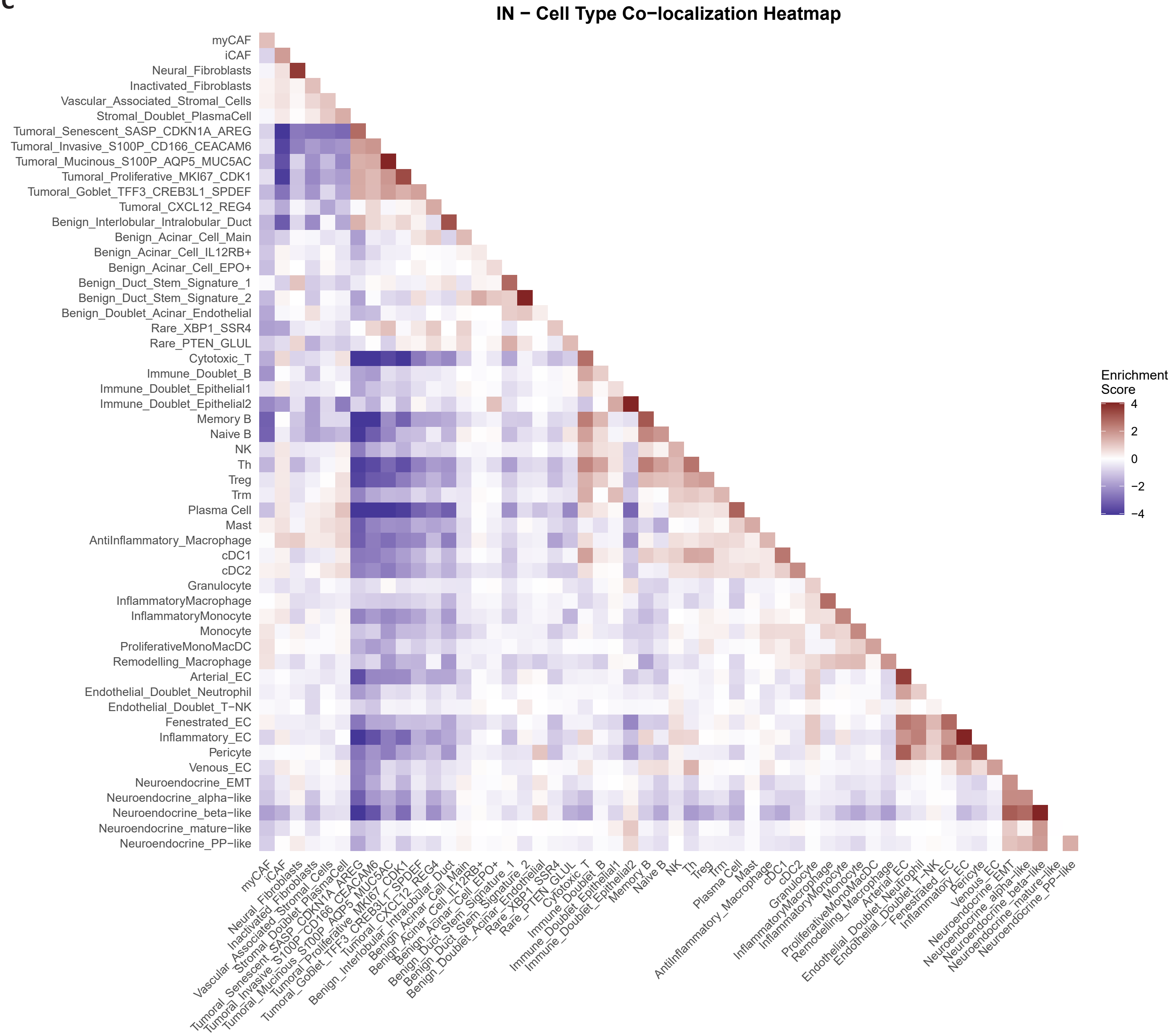

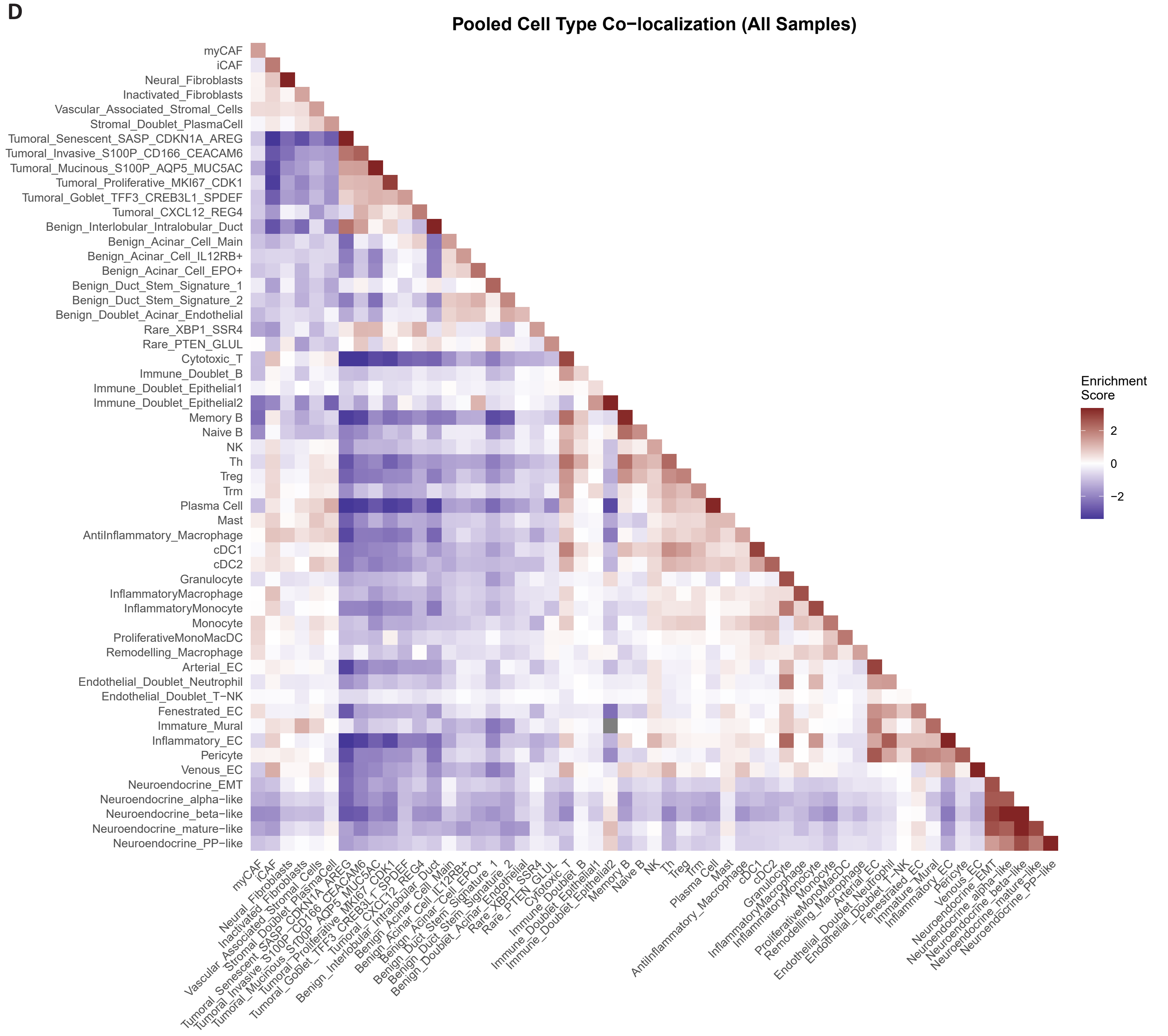

### Supplementary Fig. S8

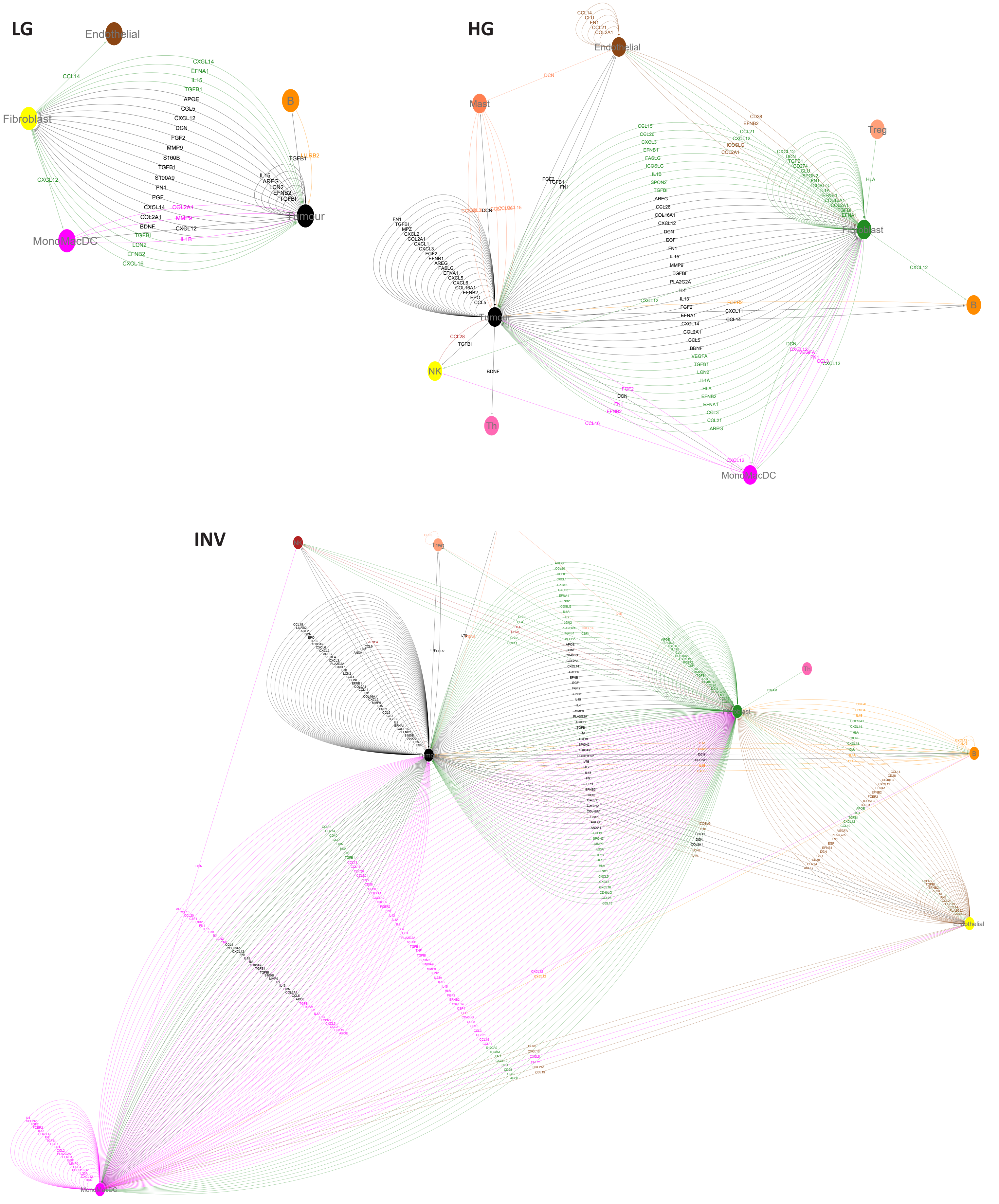
