## Supplementary Table 2 for "Spatial multi-omics resolve epithelium-fibroblast gradients and highlight NESTIN-NOTCH1-expressing subepithelial fibroblasts during human pancreatic tumorigenesis"

| Target | Antibody reference | Manufacturer/catalog | Cycle |
|---|---|---|---|
| **CD45** | Alexa Fluor® 647 anti-human CD45 Antibody | Biolegend #304056 | 1 |
| **CD4** | Human CD4 Alexa Fluor® 488-conjugated Antibody | R & D #FAB8165G | |
| **CD11b** | Recombinant Alexa Fluor® 488 Anti-CD11b antibody [EPR1344] | abcam #ab307387 | 2 |
| **CD166** | Recombinant Alexa Fluor® 488 Anti-CD166 antibody [EPR2759(2)] | abcam #ab197543 | 3 |
| **CD3** | Recombinant Alexa Fluor® 555 Anti-CD3D antibody [EP4426] | abcam #ab208514 | |
| **CD133** | CD133 (Prominin-1) Monoclonal Antibody (13A4), Alexa Fluor™ 488, eBioscience™ | Invitrogen Catalog # 53-1331-80 | 4 |
| **CXCL12** | SDF1 POLYCLONAL ANTIBODY, ALEXA FLUOR® 647 CONJUGATED | Bioss #BS-4938R-A647 | |
| **CD8** | Recombinant Alexa Fluor® 555 Anti-CD8 alpha antibody [EPR21769] (ab280863) | Abcam ab280863 | |
| **CD163** | Recombinant Alexa Fluor® 594 Anti-CD163 antibody [EPR19518] | abcam #ab282114 | 5 |
| **CD20** | Recombinant Alexa Fluor® 647 Anti-CD20 antibody [EP459Y] | abcam #ab198943 | |
| **COL1A1** | Recombinant Alexa Fluor® 647 Anti-Collagen I antibody [EPR7785] | abcam #ab280968 | 6 |
| **CD74** | CD74 Antibody (LN-2) Alexa Fluor® 594 | Santa Cruz #sc-6262 AF594 | |
| **S100P** | Recombinant PE Anti-S100P antibody [EPR6143] | abcam #ab306247 | 7 |
| **SPP1** | Recombinant Alexa Fluor® 647 Anti-Osteopontin antibody [EPR21139-316] | abcam #ab283696 | 8 |
| **CD105** | Endoglin/CD105 Antibody (A-8) Alexa Fluor® 594 | Santa Cruz #sc-376381 AF594 | |
| **CD44v9** | PE anti-human CD44 isoform 9 (CD44v9) Antibody | Biolegend #394404 | 9 |
| **CD31** | Recombinant Alexa Fluor® 647 Anti-CD31 antibody [EPR17259] | abcam #ab305210 | |
| **αSMA** | Alpha-Smooth Muscle Actin Monoclonal Antibody (1A4), Alexa Fluor™ 488, eBioscience™ | Invitrogen # 53-9760-82 | 10 |
| **FAP** | FAP Polyclonal Antibody, ALEXA FLUOR® 647 Conjugated | Bioss by Themofisher #BS-5758R-A647 | 11 |

| Slide # | Grade | Region of interest # |
|---|---|---|
| 9 | HG | 1 |
| 11 | LG | 4 |
| | HG | 4 |
| 18 | LG | 2 |
| | HG | 3 |
| 21 | HG | 21 |
| | INV | 2 |
| 22 | N | 4 |
| | LG | 15 |
| 31 | LG | 1 |
| 33 | N | 2 |
| | LG | 19 |
| 35 | LG | 8 |
| 36 | N | 1 |
| | ADM | 1 |
| | LG | 3 |
| | HG | 1 |
| 40 | LG | 7 |
| 41 | LG | 8 |
| 53 | INV | 6 |
| 55 | N | 10 |
| | INV | 15 |
| 56 | ADM | 1 |
| 65 | ADM | 7 |
| 69 | N | 3 |
| | LG | 6 |
| | HG | 1 |
