## Supplementary Table 3 for "Spatial multi-omics resolve epithelium-fibroblast gradients and highlight NESTIN-NOTCH1-expressing subepithelial fibroblasts during human pancreatic tumorigenesis"

**Xenium In Situ - Custom gene panel list:**

ACHE, AIFM2, ALCAM, AQP5, ASNS, BDNF, BRPF1, CALCA, CDH1, CEACAM5, CHAT, CHGA, CLU, COL16A1, COL2A1, CREB3L1, DCX, EFNA1, EFNB1, EFNB2, ENG, EPAS1, EPHA2, EPHA3, EPHA4, EPHB3, EPHB4, ERGIC1, FAP, FOXA2, FSTL1, GAD1, GCG, GFAP, GKN1, GLS, GLUL, HK2, HSPA1B, INS, ITGA2, ITGA3, ITGA5, ITGB3, ITGB5, KLF2, KLF4, LCN2, LDHA, LRP1, MPZ, MUC1, MUC6, MUCL3, MYC, NCK2, NES, NEUROD1, NFIB, NKX6-2, NTF3, OGDH, OLIG2, PBX1, PDGFRB, PDX1, PECAM1, PLA2G2A, POSTN, POU2F3, PPY, PROM1, PTF1A, RBP4, RCOR1, REST, RFX6, S100P, SERPINA7, SETBP1, SIN3A, SLC17A7, SLC1A5, SLC2A1, SLC7A5, SNAI1, SNAI2, SOD2, SOX10, SOX4, SOX6, SPDEF, TGFBI, TIAM1, TPH1, TWIST1, VHL, ZBTB16, ZEB1, ZEB2
