## Supplementary Table 6 for "Spatial multi-omics resolve epithelium-fibroblast gradients and highlight NESTIN-NOTCH1-expressing subepithelial fibroblasts during human pancreatic tumorigenesis"

| Item | Distributor | Cat number |
|---|---|---|
| Adenine | SIGMA | #A2786 |
| Advanced DMEM/F-12 | Life technologies - Gibco Biobar | #12634010 |
| AggreWell HT, 5pk | Stemcell tech | #200-0570 |
| Anti-Adherence Rinsing Solution | Stemcell tech | #07010 |
| Antibiotic-Antimycotic, 100X Solution | Wisent | #450-115-EL |
| B-27 Supplement (50X), serum free | Life tech-Gibco | #17504044 |
| Corning® Matrigel® Basement Membrane Matrix, Phenol Red-free | Corning | 356237 (CB-40234C) |
| DMEM, high glucose, pyruvate (4.5g/L D glucose) | Life technologies - Gibco Biobar | #11995073 |
| D-PBS 1X (w/o Ca2+, Mg2+) , sterile | Wisent | #311-425-CL |
| FBS | WISENT | lot 112740 |
| Gentamicin [50mg/ml] | WISENT | 450-135-XL |
| GLUTAMAX I, 100X | Life technologies - Gibco Biobar | #35050061 |
| Ham's F-12 Nutrient Mix | Life tech-Gibco | #11765054 |
| Hydrocortisone | SIGMA | #H4001 |
| L-Glutamine (200 mM) | Life technologies - Gibco Biobar | #25030081 |
| Nirogacestat (Notch inhibitor) (10mM) | Ambeed | A421613 |
| N-2 Supplement (100X) | Life tech-Gibco | #17502048 |
| Penicillin-Streptomycin, 100X Solution | WISENT | 450-201-EL |
| Trypsin-EDTA (0.05%), phenol red | Life tech-Gibco | #25300062 |
| Y-27632 dihydrochloride | Biotechne Tocris | #1254 |
| 37 µm Reversible Strainer, Large | Stemcell tech | #27250 |
| 96 wells plates adherentes | Sarstedt | #83.3924.500 |
| RPMI 1640 Medium, HEPES | Life tech-Gibco | #22400089 |

**Conditioned Growth Medium**

| Components | Volume (Total: 500mL) | Final Conc. | Notes |
|---|---|---|---|
| Advanced DMEM-F12 | | | |
| FBS | 10mL | 2% | |
| Antibiotic-Antimycotic, 100X Solution | 5ml | 1% | |
| L-Glutamine | 5ml | 1% | |
| Adenine (lab stock 4.8mg/ml 50mM HCL) | 2,5ml | 24ug/ml | |
| Hydrocortisone (1mg/ml in 20% EtOH in Ultrapure water) | 200ul | 0,4ug/ml | |
| Y-27632 dihydrochloride (1mM) | 50ul | 10uM | |

**Fibroblasts outgrowth media**

| Components | Volume (Total: 500mL) | Final Conc. | Notes |
|---|---|---|---|

| | | |
|---|---|---|
| DMEM, high glucose, pyruvate (4.5g/L D glucose) | | |
| FBS | 25ml | |
| Penicillin-Streptomycin, 100X Solution | 5ml | 1% |
| GLUTAMAX I, 100X | 2.5ml | |
| N-2 Supplement (100X) | 1.25ml | |
| B-27 Supplement (50X), serum free | 0,625ml | |

**Transfer Medium**

| Components | Volume (Total: 500mL) | Final Conc. | Notes |
|---|---|---|---|
| RPMI 1640 Medium, HEPES | | | |
| FBS | 50ml | 1% | |
| Antibiotic-Antimycotic, 100X Solution | 5ml | 1% | |
